# Behavioral alignment as an organizing principle in sensory coding

**DOI:** 10.64898/2026.02.04.703828

**Authors:** Shuhong Huang, Ruben Portugues, James E. Fitzgerald

**Affiliations:** Institute of Neuroscience, Technical University of Munich, Munich, Germany; Graduate School of Systemic Neurosciences, University of Munich, Munich, Germany; Munich Cluster of Systems Neurology (SyNergy), Munich, Germany; Max Planck Fellow Group - Mechanisms of Cognition, MPI Psychiatry, Munich, Germany; Bernstein Center for Computational Neuroscience, Munich, Germany; Department of Neurobiology and Behavior, Cornell University, Ithaca, NY, USA; Departments of Neurobiology, Physics and Astronomy, and Engineering Sciences and Applied Mathematics, Northwestern University, Evanston, IL, USA; NSF-Simons National Institute for Theory and Mathematics in Biology, Chicago, IL, USA

## Abstract

The ultimate goal of sensory coding is to extract and represent the cues required for adaptive motor output. This suggests that sensory codes and behavioral outcomes may align, and a variety of studies have argued that both biological and engineered sensory systems represent stimuli similarly when they play similar roles in behavior. However, the extent to which behavioral demands determine sensory coding throughout the brain is largely unknown. Here we propose that behavioral alignment is a general principle that organizes sensory representations, and we show that carefully measured behavior can predictively account for visual encoding across the entire zebrafish brain. We discover population codes that represent visual motion stimuli according to the optomotor responses elicited by them, indicating that information required for behavioral selection is explicitly encoded in sensory populations. Brain-like neuronal responses result when sensory codes are optimized for efficiency and aligned to behavior. These results provide a paradigm for understanding sensory representations through the behaviors they drive.

## 1 Introduction

Adaptive behavior requires animals to perceive stimuli and flexibly select actions based on what they sense in the environment. Sensory systems facilitate this process by encoding various features of the stimulus through patterns of neural activity. Some of these features correspond to simple physical properties of the stimulus, but more abstract features could better align with behavioral outcomes (Figure 1). However, there is no consensus on the specific features that sensory systems encode, let alone the normative and mechanistic principles that determine the encoding.

**Fig. 1.**
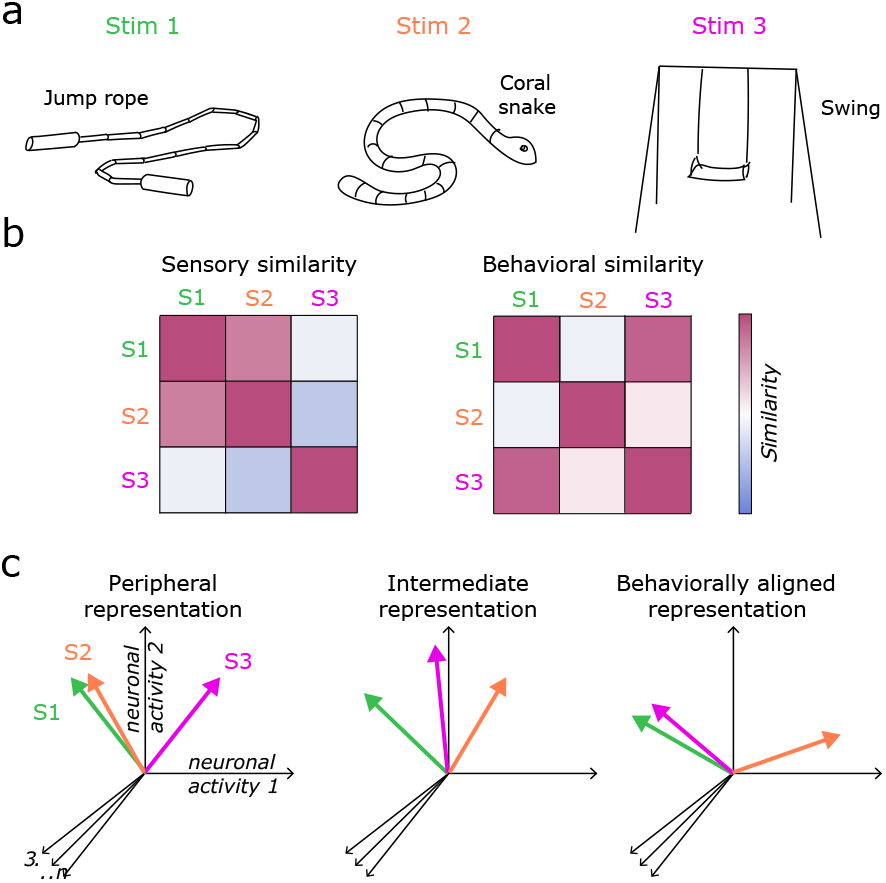
The concept of behavioral alignment. **a**, Three example visual stimuli. **b**, Matrices quantifying different notions of similarity between the three stimuli. Based on raw sensory qualities like shape and texture, Stimuli 1 and 2 are more similar to each other than Stimulus 3 (left). However, Stimuli 1 and 3 are more similar in terms of the behaviors they drive (e.g., approach and play) than Stimulus 2 (e.g., freeze and back away) (right). **c**, Possible neuronal representations of the stimuli. Neurons near the sensory periphery likely reflect physical stimulus properties, such that Stimuli 1 and 2 evoke similar population activity patterns (left). A behaviorally aligned representation instead codes stimuli according to their behavioral outcomes (right). Behaviorally aligned representations could be mechanistically caused by the stimulus but normatively predicted by the downstream behaviors. Multi-stage sensory processing may give rise to intermediate representations between these extremes (middle).

One prominent perspective is that sensory coding constructs an internal representation that is as informative as possible about the external world [1–3]. Based on this view, the efficient coding hypothesis proposes sensory codes that are optimized to minimize the redundancy of sensory encoding or maximize the mutual information between neural responses and natural stimuli. This framework has predicted sparse and decorrelated sensory codes similar to those observed in many experimental studies [4–14]. However, the successes of efficient coding largely pertain to peripheral sensory systems. This is perhaps unsurprising considering that the brain’s primary function is behavior, and sensorimotor pathways should discard sensory information when that information is not behaviorally relevant [15, 16].

An alternate perspective supposes that sensory coding is optimized to support specific behavioral tasks [1, 17–21]. This paradigm has found much recent success by predicting brain-like representations in multiple layers of sensory processing [22–29]. For example, when deep neural networks are trained using machine-learning methods to perform image classification, the hidden layer units can linearly predict neuronal data across multiple regions of the ventral stream. During the training process of these networks, hidden layers gradually extract task-relevant features from the input and task-irrelevant information is compressed [30–32], effectively aligning the representations to the behavioral output. Therefore, the brain-like representations of hidden layers appear to meet particular demands of the task.

In many ethological behaviors, such as hunting, escaping, and courtship, the task is sensorimotor, fundamentally requiring that sensory cues drive specific motor outputs [33, 34]. However, the extent to which the motor demands of ecological behavior predict sensory coding throughout the brain has not been investigated. One prominent behavior is the optomotor response (OMR), which stabilizes an animal’s position in its environment by generating locomotor actions that counteract optic flow from involuntary movements [35–40]. This innate behavior develops in the larval stage of zebrafish, wherein the fish is still translucent and small enough to permit cellular-resolution optical imaging of almost all its around 100,000 neurons [41, 42]. Moreover, neural coding of optomotor stimuli in several visual areas of the zebrafish brain has been suggested to be organized according to behavioral outcomes [43–45], a promising observation that motivates a comprehensive analysis of how sensory coding is organized at the whole-brain scale to support this critical sensorimotor behavior.

## 2 Results

### 2.1 Behavioral output and sensory codes in the optomotor response of zebrafish larvae

To understand sensory coding through its sensorimotor task, we first characterized the motor output of a sensorimotor behavior. In particular, we tracked larval zebrafish swimming during the optomotor response, a visuomotor behavior wherein fish generate locomotor actions triggered by optic flow (Figure 2a). The presented stimuli can be summarized into two categories of binocular visual motion: translational motion and rotation-like motion (Figure 2b). Those stimuli combined eye-specific monocular motion patterns, either drifting gratings or rotating windmills, into binocular ones, such that the fish could experience various stimuli and generate diverse behavioral responses.

**Fig. 2.**
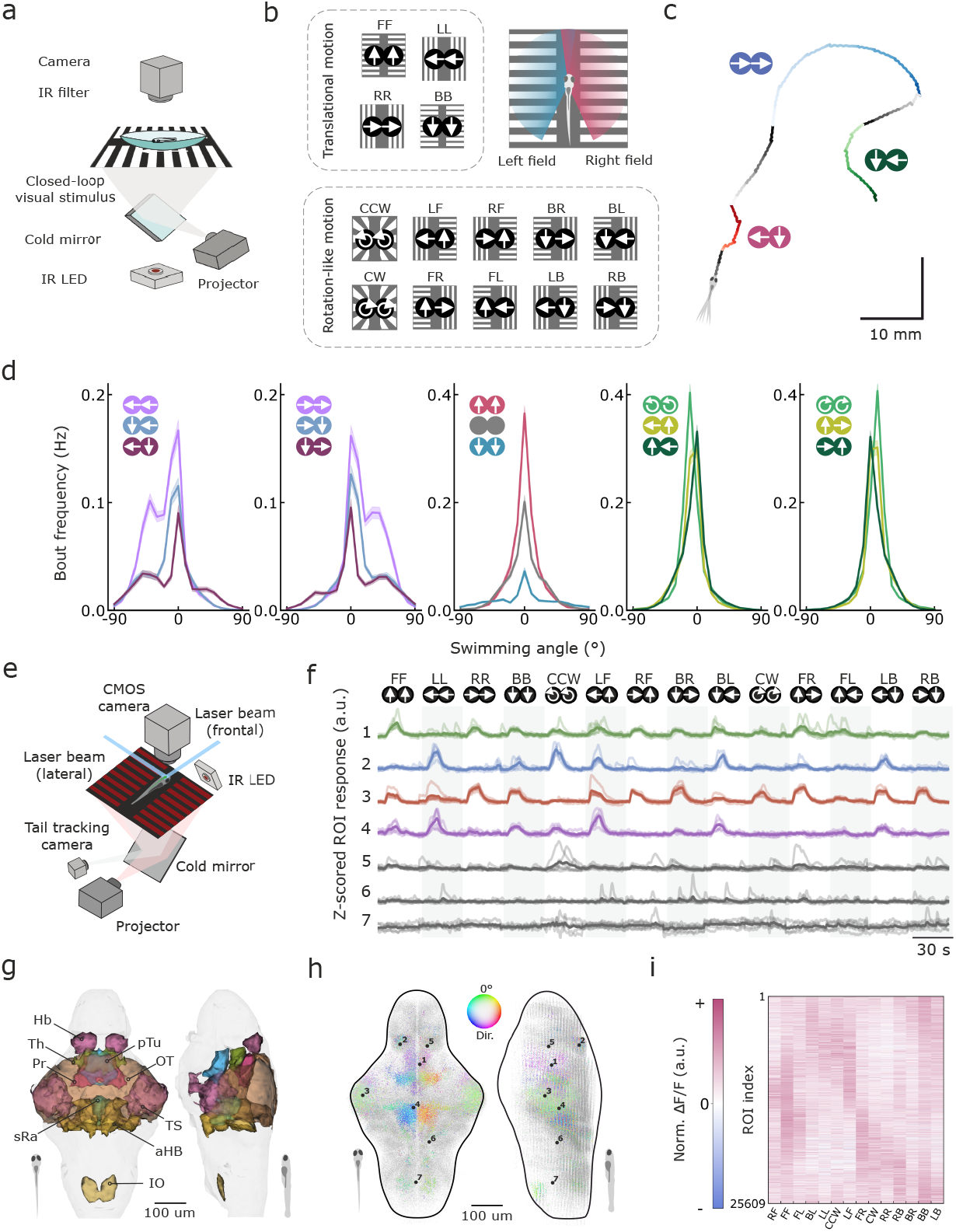
Behavioral and neural responses under visual motion stimuli. **a**, Schematic diagram of the experimental setup used for behavioral studies. **b**, The stimulus set including 14 combinations of monocular visual motion. Top left: Translational motion stimuli, where both eyes receive synchronized directional motion. Bottom: Rotation-like stimuli, each consisting of distinct monocular motion patterns. Top right: Schematic illustration of stimulus presentation. The monocular visual motion is not detected by the contralateral eye. **c**, An example swimming trace during the freely swimming experiment. **d**, Histograms of bout angle frequencies. Shaded area represents standard error of the mean (SEM), n=67. **e**, Schematic diagram of the setup used for whole-brain imaging experiments. **f**, Examples of raw (light) and average (dark) calcium traces from regions of interest (ROIs) in response to 14 visual motion stimuli. The positions of these ROIs are shown in **h. g**, Examples of reference brain regions, labeled based on [49]. Pr, pretectum; Hb, habenula; TS, torus semicircularis; OT, optic tectum; aHB, rhombomere 1 & 2; IO, inferior olive; sRa, superior raphe; pTu, posterior tuberculum. **h**, Distribution of motion-sensitive ROIs (n=25,605), color-coded by their preferred translational motion direction, alongside other detected ROIs (n=230,445) across 12 *Tg(elavl3:GCaMP6s)* fish. **i**, Δ*F/F* of all selected sensory ROIs under 14 stimuli. Neural activities are normalized for each ROI.

We quantified this sensory-induced behavior by how often the fish swam in each possible direction. For larval zebrafish, the motor output of OMR is discrete swimming events called bouts (Figure 2c). Therefore, we binned swimming directions into nineteen 10°-wide intervals and computed the bout frequency for each bin (Figure 2d). Compared to spontaneous swimming, translational visual motion elicited swimming towards the stimulus direction and suppressed that towards the opposite direction. In addition, nasal motion paired with backward motion elicited more turns facing the stimulus direction and suppressed opposite turns more effectively than temporal motion, consistent with previous findings [35]. However, this pattern was reversed when combined with forward motion, with nasal motion triggering fewer turns towards the stimulus direction and more turns towards the opposite direction compared to temporal motion. This observation shows that sensorimotor computation in OMR is not simply summing the contributions of monocular visual motion.

We identified sensory neurons as those that responded reliably across trials to at least one visual stimulus. Using whole-brain calcium imaging, we extracted regions of interest (ROIs) and their fluorescence traces from 12 fish with expressing GCaMP6s [46] in nearly all neurons (Figure 2e-h). Sensory ROIs were distributed broadly across the brain (Figure 2g,h; Extended Data Figures 1,2) and were anatomically distinct from motor-related neurons that correlated with individual bouts (Extended Data Figure 3). Consistent with previous studies, sensory ROIs in optic tectum were mostly tuned to motion along body axis, whereas those in pretectum and anterior hindbrain were more tuned to leftward or rightward motion [35, 43]. The sensory responses of all selected ROIs are summarized in Figure 2i, sorted based on their preferred directions.

### 2.2 Multiple behavioral elements explain single-neuron responses

A major goal of sensorimotor computation is to format sensory signals in a way that motor systems can read out to generate behavior [1, 17, 18]. This suggests that sensory signals close to motor output might encode sensory stimuli similarly to their impact on behavior. We thus asked whether we could interpret the sensory codes measured through the zebrafish brain in terms of the measured patterns of stimulus-evoked behavior. For instance, does each neuron’s stimulus response match a basic pattern also observed in the behavioral output?

We first learned a group of behavioral elements using sparse dictionary learning on the stimulus-triggered behavioral output (i.e. bout frequencies during stimulus subtracting those in darkness) (Figure 3a-b). This method sparsely encodes each sample with only a few learned elements and has been used successfully to predict receptive fields of macaque V1 neurons from natural images [7]. The reconstruction error of behavioral output decreased as more elements were added (Extended Data Figure 4a), with six elements substantially reducing the error (see Methods). To understand the extracted behavioral elements, we examined each element’s weight profile over presented visual stimuli (Figure 3c). These elements had interpretable structure across the stimulus set: left vs. right motion (1 and 6), translational vs. rotational motion (2 and 5), clockwise vs. counterclockwise rotation (3), and backward vs. lateral motion (4). In addition, we computed the cosine similarity between each behavioral element and the direction-specific bout frequencies (Figure 3d). This revealed that these elements are selective for leftward versus rightward turns or for forward swimming versus turning from a motor perspective.

**Fig. 3.**
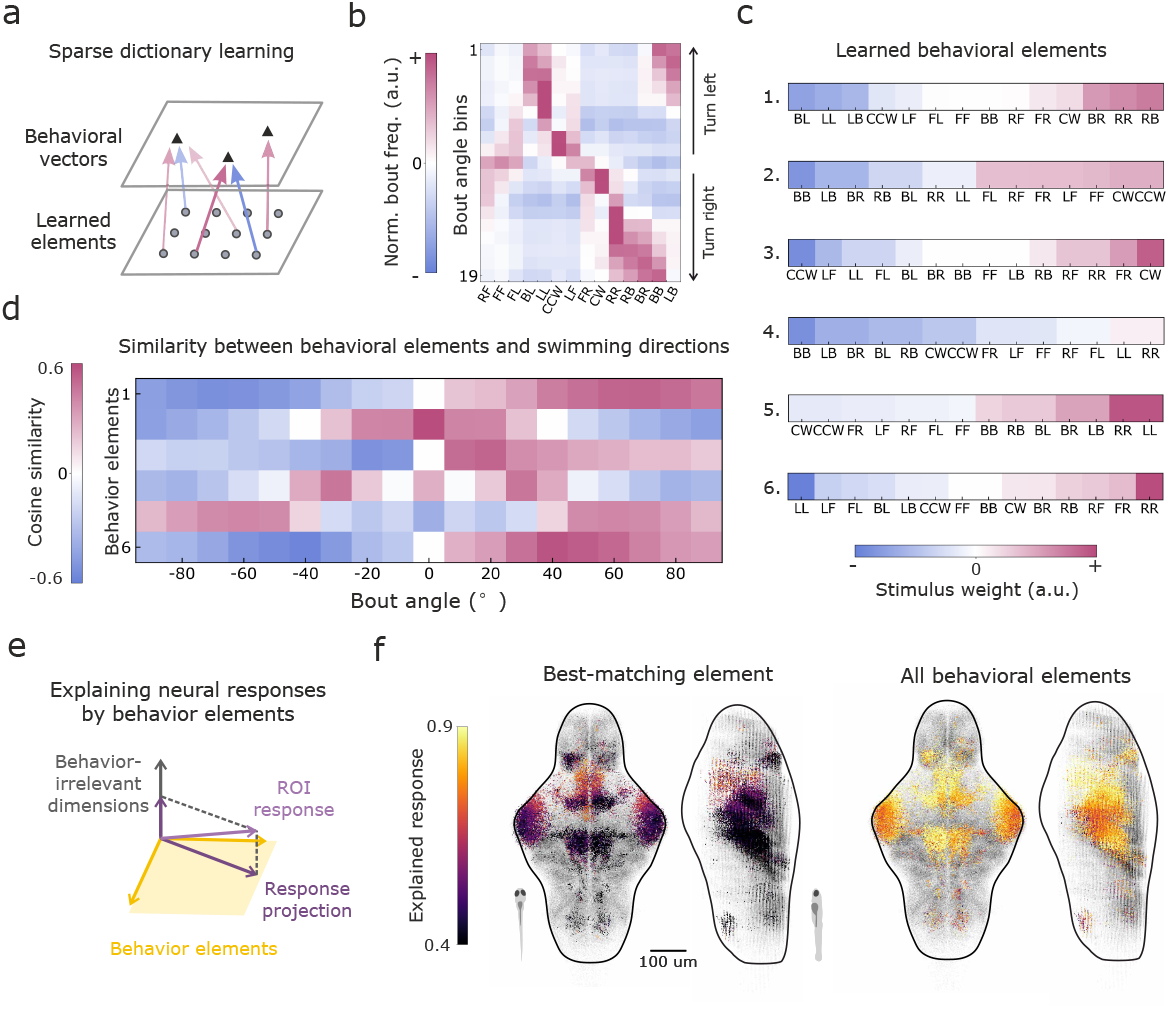
Explaining sensory responses using behavioral elements. **a**, Illustration diagram for sparse dictionary learning. **b**, Stimulus-triggered behavioral responses, with each column showing the relative bout frequency (stimulus-induced minus spontaneous swimming in darkness) in 10°bins from -95°to 95°, and the bout frequency is normalized for each angle bin. **c**, Weights of learned behavioral elements for each stimulus. Stimulus order is sorted by its weight in each behavioral element. **d**, Cosine similarity between each behavior element and bout frequencies within each bout angle bin in **b. e**, Illustration diagram of decomposing ROI responses using behavioral elements. **f**, Explained response of each ROI by best-matching behavior elements (left) and by the space spanned by all six behavioral elements (right).

Most ROIs needed multiple behavioral elements to fully explain their activity. To see this, we first tested whether single ROI responses were aligned with a single behavioral element. For each ROI, we projected its response vector onto each behavioral element (Figure 3e). The component of the sensory response along each behavioral element quantified how much of the sensory response could be explained by the behavioral element. We found that a single behavioral element was usually not enough to explain single-ROI responses (Figure 3f). However, the set of behavioral elements as a whole could accurately reconstruct these single-ROI responses (Figure 3g; Extended Data Figure 4). Thus, the response level of a single ROI reflects a combination of multiple behavioral elements, rather than uniquely representing one element. A related analysis based upon principal components analysis produced similar results (Extended Data Figure 4), but the behavioral basis functions produced by principal component analysis were less interpretable and less aligned to single ROI activity.

### 2.3 Behavior-aligned stimulus representations in sensory populations

Having shown that the behavioral elements largely explain sensory activity, we next asked a stronger question: are stimulus representations aligned between sensory populations and behavioral output? If sensory codes align with behavior, then populations of sensory neurons should represent stimuli similarly when they produce similar behaviors [47]. To illustrate this point, we first considered neural network models where raw sensory inputs are transformed into behavioral outputs through multiple feedforward layers of neural representation. We focused on deep linear neural networks (i.e., multilayer perceptrons wherein all activations are the identity function), because the representational structure of each hidden layer can be understood comprehensively using analytical methods [48] (see Methods). When we trained a deep linear network to transform random sensory cues into the behavioral responses (Figure 4b), we found that multiple layers of “sensorimotor” processing gradually produced neuronal representations with stimulus correlations that match the behavior (Figure 4c), as expected from theory. This result did not depend on the details of the sensory representation, and modeling direction-selective retinal ganglion cells as the network input produced similar results (Extended Data Figure 5).

**Fig. 4.**
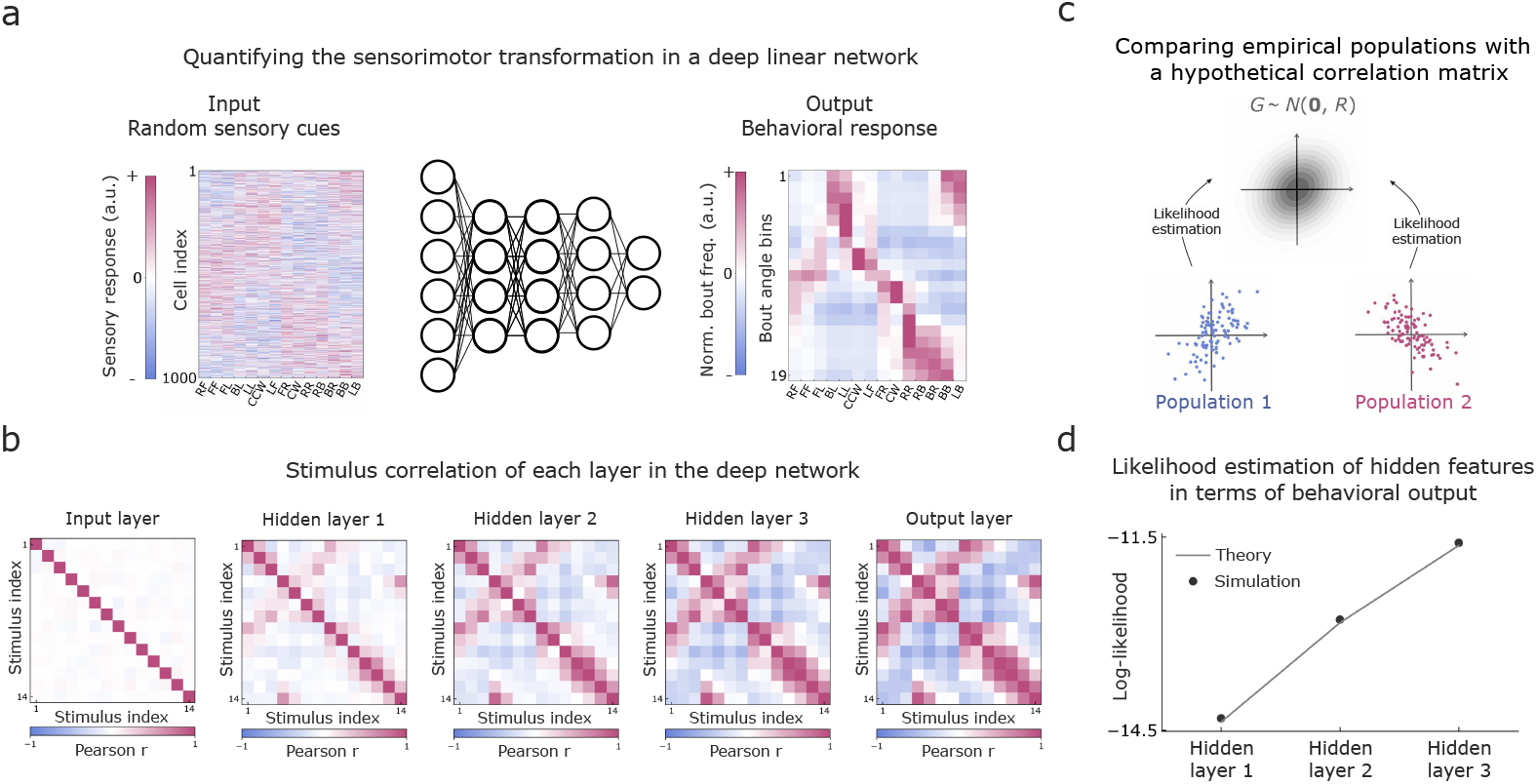
A likelihood-based method to quantify behavioral alignment. **a**, A deep linear network transforms randomly sampled sensory responses into behavioral responses. **b**, Stimulus similarity matrix of each layer in the deep linear network for the random sensory cues in panel **a. c**, We quantify the behavioral alignment of neural populations by how computed likely the neuronal responses are given a Gaussian model with the behavioral correlation matrix. **d**, Likelihood estimation of features in hidden layers in terms of stimulus similarity matrix of behavioral output. Error bar represents SEM for 20 independently sampled random sensory cues.

To explore whether this concept can explain the experimental data, we developed a quantitative framework to determine if the pair-wise stimulus correlations of sensory population codes match behavior. This framework is similar to representational similarity analysis [47], but it differs by quantifying the alignment between a given correlation matrix and the observed data using the log-likelihood of the data under a Gaussian model with zero mean and covariance defined by the specified correlation matrix (Figure 4a, Methods). For instance, if we define the Gaussian distribution with stimulus correlations from the behavioral response, then a sensory population with high likelihood would exhibit similar neural responses to stimuli that elicit similar behavioral responses, and vice versa. This framework successfully captured the layer-by-layer transformations observed in our deep linear neural network model, with deeper layers exhibiting higher log-likelihood as their correlation matrices became aligned to the output (Figure 4d).

We therefore proceeded to apply our method to the experimental data. As above, we used the stimulus correlation matrix of the behavioral responses (Figure 5a) to define the Gaussian distribution in the proposed framework. We then estimated likelihoods for sensory populations across twelve anatomical brain regions that contain the most sensory neurons [49]. Among these regions, the estimated likelihood was significantly higher in the pretectum, habenula, tegmentum, and the reticular formation in rhombomeres 1-2 (Figure 5b, Extended Data Table 1). Sensory populations in the hindbrain showed especially strong behavioral alignment in terms of the estimated likelihood. Visualizing the stimulus correlation matrices also supported these findings, as these five regions exhibit stimulus correlation patterns closely matching those of the behavioral responses (Figure 5c,d). In contrast, the optic tectum and torus semicircularis exhibited strong monocular patterns, while the thalamus, hypothalamus, and posterior tuberculum did not display clearly interpretable patterns (Extended Data Figure 6). These sensory populations still encode visual motion stimuli and may support other behaviors that are guided by optic flow information. Overall, these results indicate that specific sensory populations encode behavioral responses, and that similarities in behavior are encoded in similarities in sensory responses.

**Fig. 5.**
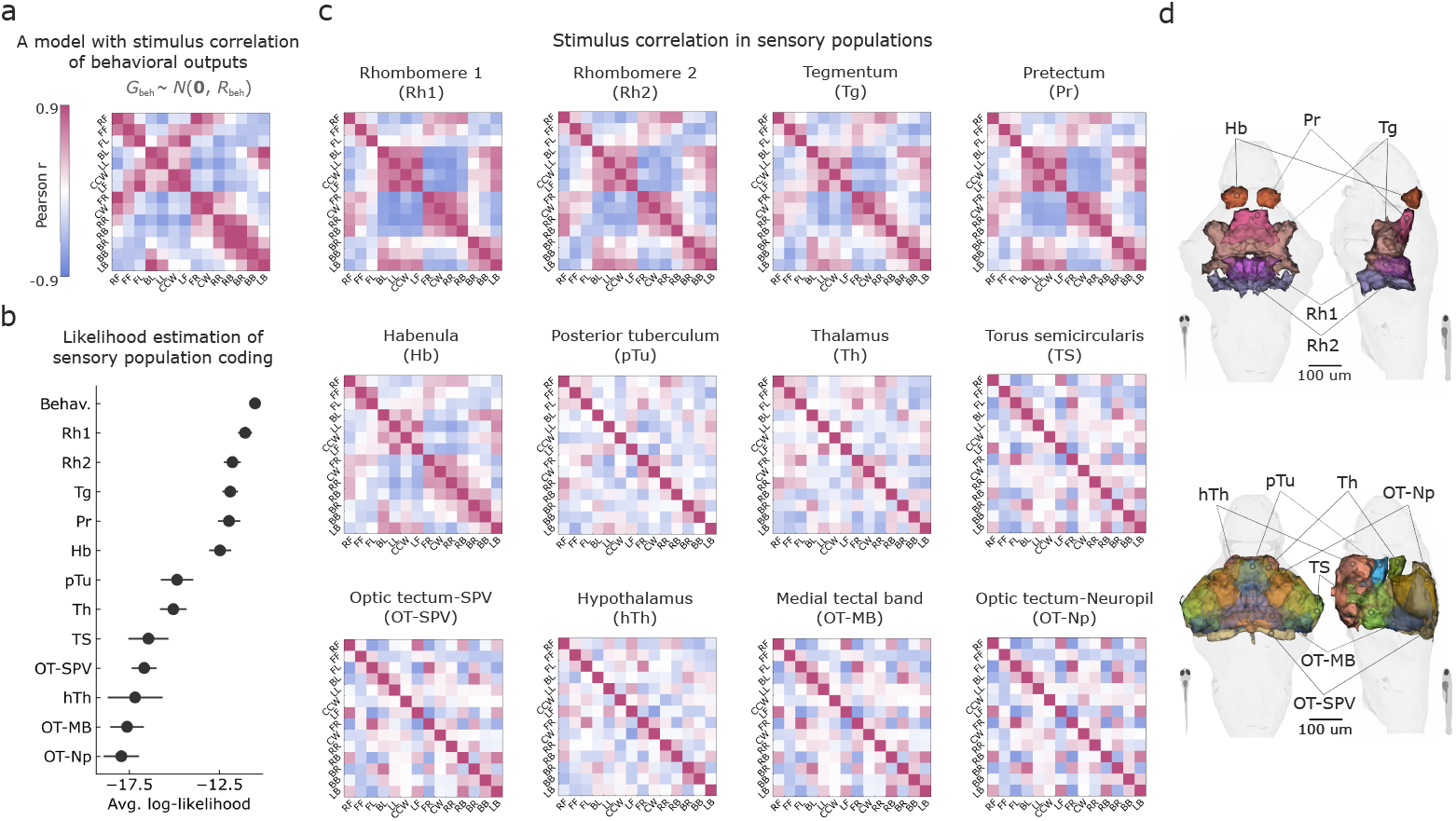
Stimulus similarity measurement of anatomical sensory populations. **a**, Stimulus similarity of behavioral output. **b**, Likelihood estimation of anatomical sensory populations from the model with stimulus similarity of behavioral output. Error bar represents SEM among the imaged fish. **c**, Stimulus similarity matrices of anatomical sensory populations. **d**, Reference of brain anatomy. Top: Pr, pretectum; Hb, habenula; Tg, Tegmentum; Rh1, rhombomere 1; Rh2, rhombomere 2. Bottom: TS, torus semicircularis; TH, thalamus; hTh, hypothalamus; pTu, posterior tuberculum; OT-Np, optic tectum - neuropil; OT-MB, optic tectum - medial band; OT-SPV, optic tectum - stratum periventriculare.

### 2.4 Efficient coding of stimuli driving behavior

We next asked what principles are needed to predict single ROI responses. Given that sensory population codes exhibit stimulus correlations similar to the behavioral output, we assumed that the neural activity could be transformed into the behavioral responses through a linear readout (Figure 6a). Moreover, we modeled the readout weight matrix as random and Gaussian to preserve the stimulus correlations across the transformation (Supplementary information). Many representations are consistent with these assumptions, and, following the many successes of efficient coding theory, we hypothesized that biology would use an efficient representation that minimizes the population activity over neurons and stimuli (i.e. minimizes the Frobenius norm, Figure 6b_1_, Methods).

**Fig. 6.**
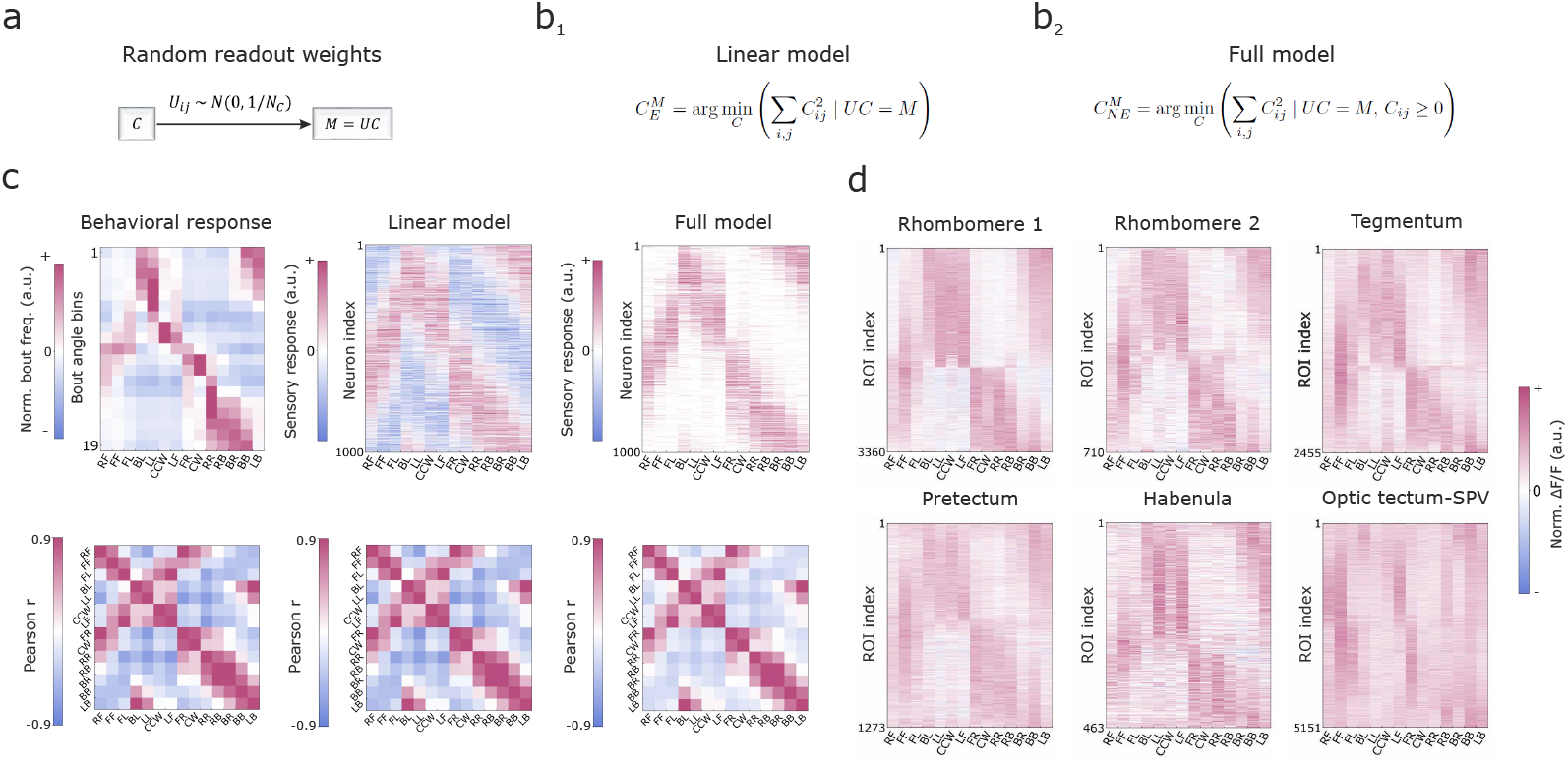
A simple model for efficient coding of stimuli driving behavior generates brain-like sensory activities. **a**, Illustration of modeled neural activities being read out into behavioral outputs through a Gaussian connection matrix. **b**, Two optimization models to generate neural activities in terms of behavioral output **b**_**1**_, Neural activities with minimal *L*_2_-norm; **b**_**2**_, Non-negative neural activities with minimal *L*_2_-norm. **c**, Sensory activities (top) and stimulus similarity matrix (bottom) for behavioral response, prediction of the linear model, and prediction of the full nonlinear model. **d**, Neural activities of six anatomical sensory populations.

These minimal assumptions were insufficient to explain the single-ROI activity patterns measured experimentally. Figure 6c (left and middle columns) compares the behavioral responses and the modeled sensory activity. Consistent with our theoretical expectations (Supplementary information), these two representations were closely aligned and shared the same stimulus correlation matrix. However, the modeled sensory activities were dense and symmetrically distributed across positive and negative values, which contrasts with the sparse and non-negative characteristics of actual neural responses (Figure 6d).

However, a modified model that additionally imposed a non-negativity constraint was able to account for the single-ROI activity patterns (Figure 6b_2_). Similar to the previous model, we both solved this model analytically and generated sensory activity numerically (Supplementary information). Although this model’s stimulus correlation matrix no longer perfectly matched that of behavior, this was also true of the experimental data, and many structural patterns were preserved (Figure 6c, right column). Comparing the modeled sensory activity to experimental data, the modeled activity captured many salient patterns, such as anti-correlated activities between left and right stimuli, forward and backward stimuli, and clockwise and counterclockwise stimuli (Figure 6d, Extended Data Figure 8). Moreover, the modeled activity was sparse and non-negative. The proposed model therefore generates brain-like single-neuron sensory activity based only on behavioral responses, suggesting that sensory representations can be understood in term of their behavioral alignment.

## 3 Discussion

The present study demonstrated that the zebrafish visual system encodes stimuli in a manner aligned with the behavioral responses they elicit. Our study provides a conceptual framework for understanding recent work suggesting that sensory neurons in the zebrafish respond to physically diverse stimuli that trigger similar directed locomotion [35, 43, 44, 50]. It also provides a new perspective on the widespread encoding of valence in diverse brains [51–53], as attractive stimuli drive approach behaviors and aversive stimuli drive avoidance behaviors. How do animals benefit from these behavior-aligned sensory codes? One potential benefit is to avoid mistakes in action selection. Since stimuli that drive distinct actions are well separated as sensory codes, even when environmental or internal noise distorts sensory representations, the noise is unlikely to result in large behavioral differences.

Behavior-aligned sensory coding may provide a new normative principle of sensory encoding. Here we presented a proof-of-concept optimization model that could predict single-neuron activity patterns by hypothesizing that the population code is efficient, non-negative, and aligned to behavior (Figure 6). This normative model was not fit to neural data, suggesting that the model captures critical behavioral objectives and mechanistic constraints shaping neural computation. Previous normative models of sensory coding based on efficient coding predict that sensory codes should adapt to changing sensory statistics of the animal’s environment [54–57]. The behavior-aligned coding principle likewise suggests that sensory representations should adapt to the changing statistics of behavior. For example, during training on perceptual decision-making tasks, sensory representations of task-relevant stimuli changed with behavioral performance [58–60], and such changes did not arise from passive exposure to the stimuli alone [61, 62]. These results might reflect the alignment of learning-shaped sensory representations and stimulus-driven behavior.

Our proposed method to quantify the behavioral alignment of sensory population codes could be extended in important ways. Our method utilized the behavioral correlation among the stimuli to model a probability distribution of behavior-aligned representations and assessed population codes based on their likelihood. This likelihood measurement conceptualizes the observed neural activity as samples from an explicit generative model, and the resulting comparisons are closely related to representational similarity analysis (RSA) [47, 63]. For instance, our method measures the alignment of global structures, as also captured by RSA [64–66]. However, our likelihood-based framework also allows us to consider alternate generative models and could enable flexible extensions to model alternative response distributions and noise correlation between neurons. For example, Fig. 6 shows a non-Gaussian generative model that explained single neuron activity better than its Gaussian alternative, and it would be interesting to extend our method to this case and future improved models.

Although this study focused on a simple sensorimotor behavior, the proposed framework has the potential to extend to more complex and flexible behaviors. While the importance of behavior in shaping sensory codes has been recognized previously [19–22], we went further by carefully measuring the animal behavior to explain sensory activity based upon it. Moreover, we found that coarse behavioral measures could obscure important details of the code (Extended Data Figures 9, 10). Going forward, recent advances in behavioral quantification could allow for richer descriptions of behavior across the animal’s natural behavior repertoire [67–69], and the motor commands generating it might provide a particularly insightful representation that quantifies how ecological motor behaviors are read out from sensory codes [70]. We therefore anticipate that more comprehensive investigations of animal behavior and motor control will deepen our understanding of sensory systems, and future work should map sensorimotor circuits based upon the gradual or abrupt emergence of behaviorally aligned representations across the the cell types and brain regions that transform sensation into behavior.

## Acknowledgments

We would like to thank Haim Sompolinsky and Dan Lee for early input on the model. SH is funded by the European Union’s Horizon 2020 research and innovation programme under the Marie Sklodowska-Curie grant agreement #813457. SH and RP were funded by the German Research Foundation (DFG) under Germany’s Excellence Strategy within the framework of the Munich Cluster for Systems Neurology (EXC 2145 SyNergy, identifier 390857198). RP was supported by start-up funds from Cornell University. JEF acknowledges support from the Howard Hughes Medical Institute and the National Institute for Theory and Mathematics in Biology through the National Science Foundation (grant number DMS-2235451) and the Simons Foundation (grant number MPTMPS-00005320).

## 4 Methods

### 4.1 Animals

All experiments were performed with 6–8 days post-fertilization (dpf) zebrafish larvae (*Danio rerio*) of yet undetermined sex. The Tüpfel long fin (TL) wild-type strain was used for freely swimming behavioral experiments. The nacre (mitfa-/-) [71] transgenic zebrafish line Tg(elavl3:GCaMP6s+/+) [72] was used for functional imaging experiments. All procedures related to animal handling were conducted in accordance with protocols approved by the Technische Universität München and the Regierung von Oberbayern.

### 4.2 Experimental details

#### 4.2.1 Free-swimming behavioral experiments

Fish were placed in a 15 cm shallow glass dish (Duran group, watch glass, nr. 216-0281) on a clear acrylic support covered with a diffusive screen and illuminated from below using an array of IR LEDs. Freely swimming larvae were monitored using a high-speed camera (Ximea) running at 180-200 fps, equipped with a 8 mm lens (Edmund Optics) and an 830 nm long pass filter (Edmund Optics). The visual stimuli were displayed from below using a microprojector (Asus P2E or optoma 1050ST) and a cold mirror (Edmund Optics). An open-source Python software, Stytra [73], was used to track fish swim dynamics and to generate the closed-loop optic flow motion. The visual stimuli were presented below on a 10 cm by 10 cm screen, covering the 10 cm diameter circle of the dish that was filled with water. The grating patterns were either parallel or perpendicular to the body axis of fish, depending on the stimulus, and moved at 10 mm per second with the spatial period of 1 cm. The windmill patterns were centered on the fish fish, had 12 arms and moved at 36 degrees per second. To independently stimulate each eye, a 1.25-cm-width black bar was presented below the fish. The whole set of visual stimuli was presented 40 repetitions, in a different random order each repetition. Each trial of one stimulus contained 5 s of a static pattern and 10 s of motion. During the presentation of all visual stimuli, 3-min pauses were added (after 10, 20, 30 repetitions) to record spontaneous swimming. The extracted position and orientation were further analyzed with custom-written software in Python.

#### 4.2.2 Whole-brain imaging experiments

Animals were embedded in 2.2% low-melting-point agarose (Thermo Fisher) in a customized lightsheet chamber with two glass coverslips sealed on the side and a square of transparent acrylic on the bottom for behavioral tracking. The chamber was filled with water from the fish facility system and agarose around the tail was removed to enable unconstrained tail motion. To record whole-brain neural activity, a custom-built lightsheet microscope was used [74]. A 488 nm wavelength laser source (modulated laser diodes; Cobolt) was used to produce a pair of approximately 1.5-mm laser beams with two Galvo scanners (Thorlab) at 500 Hz. The two laser beams were separately conveyed through the glass windows on the fish chamber. A custom-written Python software [75] (https://github.com/portugueslab/sashimi) was used to coordinate the position of two laser beams and the collection objective. The resulting imaging data had a resolution of 0.6 × 0.6 *µ*m per pixel and 8 - 9 *µ*m z-axis gap for 21 planes at 2 Hz. The visual stimuli were presented through a microprojector (Optoma 1050ST) and a red long-pass filter (Kodak Wratten 25) to avoid contamination of the green fluorescence. The 14 stimuli were repeated 6 times in randomized order in each repetition. Each trial of one stimulus contained 21.5 s of a static pattern and 8.5 s of motion to ensure that the fluorescence returned back to baseline in between trials.

### 4.3 Data analysis

#### 4.3.1 ROI segmentation

ROIs and their fluorescence traces were extracted from the raw 4D image data using the Suite2P package [76]. A detailed introduction of the algorithms used in this package can be found in [76, 77]. Compared to the default setting, we made the following changes to make it suitable for one-photon lightsheet imaging data – subpixel: 5; max overlap: 0.95; high pass: 60; nbinned: 4000; sparse mode: False; diameter: 6. After the initial ROI extraction, we did not use the morphology-based ROI filtering in Suite2p. Instead, we discarded the ROIs with their aspect ratios larger than 1.25 and with their sizes larger than 350 pixels or smaller than 50 pixels. In addition, all ROIs outside the brain were removed. These steps helped remove non-cell ROIs that were artifactually generated. The raw traces of ROIs were preprocessed by a linear detrending to compensate drifting and a low-pass filter of 0.5 Hz to remove the scanning laser beam artifact.

#### 4.3.2 Brain registration

All images were registered to the Zebrafish Brain Browser reference [49] using an open-source Python package, ANTsPy [78] (https://github.com/ANTsX/ANTsPy). The registration included three steps. First, we manually labeled around 30 points in both brain stacks to compute an initial affine transformation. The labeled points were selected across the brain and around clear structures such as the border between midbrain and hindbrain, the anterior commissure in sub-pallium, and the shell of interpeduncular nucleus. We then used the `TRSAA` method to perform an automatic affine transformation for refinement. Two parameters were changed from the default setting – aff_sampling: 5; flow_sigma:0. Finally we used the `SyNOnly` method to perform diffeomorphic transformation with the following parameter changes – syn_samping: 20; flow_sigma: 1; reg_iterations: [20,15,10]. The brain region labels were also adopted from the anatomical atlas of [49].

#### 4.3.3 Whole-brain screening of sensory neurons

The neurons with the most consistent responses to visual stimuli were designated as sensory neurons, calculated as follows. For each ROI, we initially reconstructed the neuronal fluorescence trace for each stimulus using the time period of 5 seconds before and 25 seconds after the start of visual motion. Consequently, the length of a concatenated trace was 360 frames, including 60 frames for each of the six repetitions. We then assessed whether ROIs consistently encoded the stimulus across trials via the correlation between each ROI and a modeled regressor [79]. The regressor was initialized as a basic binary trace with value 1 when visual motion was on and 0 otherwise. Then this binary trace was convolved with an exponentially decaying kernel. The time constant of this kernel should reflect the effective dynamics of GCaMP (2.6 seconds) [46] and the neuron’s intrinsic dynamics. Due to the unknown intrinsic dynamics, we computed, for each ROI, the correlation with a modeled regressor generated using time constants ranging from 2.6 to 12.6 s in 1 s steps (Extended Data Figure 1), and we considered the Pearson correlation coefficient with maximal absolute value among the time constants as the response consistency of one stimulus. A sensory ROI might respond to one or multiple stimuli, so we used the maximal response consistency among all stimuli as a ROI’s response consistency to screen sensory ROIs from all detected ROIs. To avoid over- or under-sampling any individual fish, the threshold for response consistency was set separately for each fish such that the top 10% of ROIs with highest response consistency were selected as sensory ROIs for each fish.

For the selected sensory ROIs, the median Δ*F/F* over stimulus repetitions was used to describe the strength of the response to each stimulus. In particular, for each trial, the baseline fluorescence *F* was defined as the average of the 5 seconds before motion onset, and Δ*F* was defined by the maximal absolute change from the baseline. In particular, we computed Δ*F* as the mean fluorescence of the five highest (activation) or five lowest (suppression) frames during the stimulus, minus the baseline fluorescence *F* . ROIs were classified as activated or suppressed according to whether the peak positive or peak negative deviation from baseline had the larger absolute amplitude. The final Δ*F/F* was the median value of all six trials. The 14-d stimulus response vector for each ROI was also normalized by its *L*_2_ norm to equalize the weight of ROIs in the population analysis.

#### 4.3.4 Behavioral quantification

Since the swimming behavior of larval zebrafish can be separated into discrete swimming bouts, we used an open-source Python package [80] (https://github.com/portugueslab/bouter), to extract the bouts based on the swimming velocity. The bouts extracted from the freely swimming behavior were categorized into nineteen uniformly spaced direction categories from -95° to 95° . Therefore, the bout frequency was computed as the number of bouts within each 10° bin divided by the motion duration of each visual stimulus, with both bouts and motion periods discarded when the fish was within 1 cm of the dish border.

The behavioral response matrix, *M*, was defined by the bout directions (19 rows) under 14 stimuli (14 columns). We first preprocessed the raw bout-frequency data to reduce noise. For the self-symmetric stimuli (FF, BB, dark), the bout frequency was averaged over each pair of symmetric turning directions (e.g., -10° and 10°). Except the self-symmetric stimuli, for a pair of mirrored stimuli (e.g., FR and LF, LL and RR), the bout frequency was averaged over each pair of symmetric turning directions (e.g. -10° in FR/LL and 10° in LF/RR). We then subtracted the spontaneous bout frequency from each column in the behavioral matrix to obtain the stimulus-triggered behavior. Finally, we normalized the data for each bout direction by its *L*_2_ norm. The last step was performed for two reasons: (1) a large-angle bout shifts the body direction much more than a small-angle bout, so the same bout frequencies lead to distinct behavioral consequences across bout directions; (2) the baseline bout frequencies in spontaneous swimming are also distinct across bout directions, so an overall increase of bout frequency leads to different gains for each bout direction.

### 4.4 Quantitative modeling

#### 4.4.1 Explaining sensory responses by behavioral elements

We used the `DictionaryLearning` method from the `sklearn` Python package to extract behavioral elements from the behavior matrix, *M* . This method seeks a group of vectors that could linearly construct the behavior matrix with *L*_1_ regularization that leads to sparse weights,

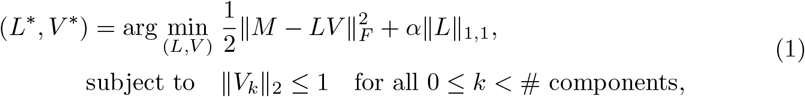

where *L* represents the weight matrix, *V* and *V*_*k*_ represent the matrix of learned behavioral elements and the *k*-th behavioral element, ∥ · ∥_*F*_ represents the Frobenius norm, ∥ · ∥_1,1_ represents the entry-wise *L*_1_ norm, ∥ · ∥_2_ represents the *L*_2_ norm, and *α* is the regularization strength. We performed a grid search over *α* from 1, 0.1, 0.01, 0.001 and found that the regularization parameter *α* = 0.1 provided a good trade-off between reconstruction error and sparsity. The relative reconstruction error was computed as

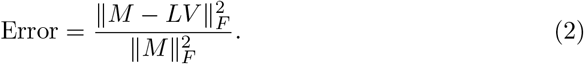

The number of behavioral elements was selected based on whether adding an additional component substantially reduced the reconstruction error. In particular, the reductions in error from adding the 2nd to the 14th components were 0.2770, 0.0828, 0.0718, 0.0500, 0.0347, 0.0154, 0.0117, 0.0160, 0.0039, 0.0025, 0.0004, 0.0009, and 0.0056. Because adding each of the first 6 elements reduced the error much more than adding any subsequent element, we extracted 6 behavioral elements. After obtaining the behavioral elements, we computed the cosine similarity between each behavioral element and bout frequency within each angle bin to assess the role of the behavioral elements in constructing the behavior matrix.

Both sensory ROI responses (**r**) and behavioral elements (**b**) can be considered as vectors in a 14-d stimulus space. We computed the explained response (ER) of one ROI and one behavioral element by

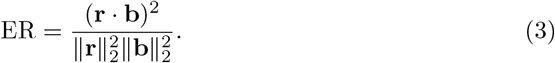

This explained response measures how much ROI responses lie in the direction of a behavioral element, and it is mathematically equal to the square of the cosine similarities between ROIs and the behavioral element. The explained response can be extended to a subspace spanned by several behavior elements. The behavior elements are noted as *B*, with 6 column vectors that represent learned behavioral elements. Since behavioral elements are not orthogonal, we extracted a group of orthogonal basis using singular value decomposition:

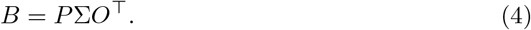

where *P* and *O*^⊤^ are orthogonal matrices and Σ is a rectangular diagonal matrix of singular values. The orthogonal basis for behavioral elements are the first 6 columns of *P*, notated as *P*_*i*_.The explained responses of all behavioral elements can be computed

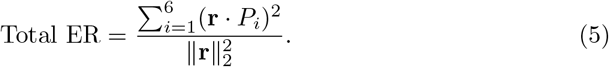

Similarly, we also computed the principal components of the behavior matrix using singular value decomposition (Extended Data Figure 4). We selected the first 7 behavioral PCs based on their explained variance. The computation of explained response was the same for behavioral PCs and raw bout frequency of each angle bins as well as for behavioral elements.

#### 4.4.2 Modeling sensory input for network training

Three types of sensory input were used in training linear deep networks to predict the motor output. For the random sensory input case, the response to each stimulus was defined as a 1000-dimensional vector independently sampled from a standard Gaussian distribution.

The other two cases relied on modeled direction-selective retinal ganglion cells. In both cases, the receptive field of each cell was one of sixteen blocks that uniformly covered the whole visual field (Extended Data Figure 5a). Within each receptive field, there were 18 modeled retinal ganglion cells. For the uniform case, these neurons were uniformly tuned to angles from -180° to 180° (Extended Data Figure 5b_1_), with tuning curves modeled by a max-min normalized von Mises probability density curve,

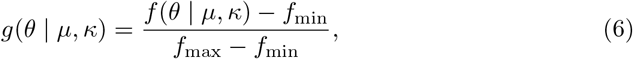

where *f* (*θ* | *µ, κ*) is the standard von Mises density curve,

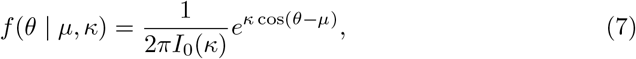

*f*_max_ and *f*_min_ are the maximal and minimal value of *f* (*θ* | *µ, κ*), *θ* is the angle of the stimulus, *µ* is the preferred tuning angle, *κ* = 0.56 is the concentration parameter (similar to a Gaussian curve with half-width at half-maximum of 90°), and *I*_0_(*κ*) is the modified Bessel function of the first kind of order 0. For the non-uniform case, *κ* was the same as above, but the preferred tuning angles were sampled based on the histogram of preferred tuning angles of voxels of retinal ganglion cell axons reported in [81]. In particular, the 18 preferred tuning angles split the areas of the histogram equally into 19 pieces. Since the reported data were recorded from one eye, we flipped the preferred tuning angle for the other eye around 0° (Extended Data Figure 5b_2_). Small independent Gaussian noise (*σ* = 0.035) was added to the modeled retinal responses to avoid zero singular values.

#### 4.4.3 Neural network training

We trained deep linear networks with three hidden layers to transform the modeled sensory codes (*H*_0_) into the measured behavioral output (*H*_4_),

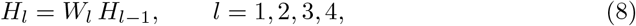

where *H*_*l*_ is a matrix that represents the network features for 14 stimuli of the *l*-th layer, and *W*_*l*_ is the weight matrix between the *l*-th layer and its previous layer. All tasks for different sensory inputs shared the same hidden architecture with *N*_*l*_ = 512 units in each of the three (*l* = 1, 2, 3) linear hidden layers. We used the Pytorch library [82] and the Adam optimizer (learning rate: 0.0001) [83] to minimize the squared error between the network output and the behavioral output,

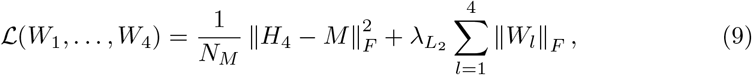

where *H*_4_ is the network output, *M* is the behavior matrix, *N*_*M*_ is the number of elements of *M* (266 in this paper), 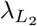 is the regularization coefficient 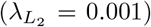, and 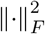 denotes the squared Frobenius norm. Before training, we performed z-score normalization to both the network inputs and outputs for each stimulus. The weight between *l*-th and (*l* + 1)-th layer of the network was initialized from a uniform distribution with the range 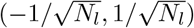. We trained the network for 20,000 iterations for the random input and 50,000 for modeled retinal ganglion cell inputs. Under this setting, the loss in Eq. 9 was 0 and the norm of weights were converged.

#### 4.4.4 Analytical expression for the hidden layer representations

Following the backpropagating kernel renormalization (BPKR) framework for deep linear networks with multiple outputs [48], we computed the expected layerwise covariance matrix of the *l*-th hidden layer as

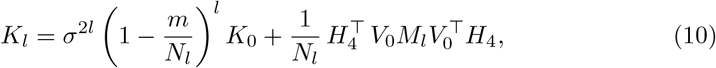

where *m* is the output dimensionality (19 swimming directions in this paper), *σ* is the hyperparameter of weight scale in the BPKR framework (*σ* = 0.1 in this paper), and the input kernel is defined as 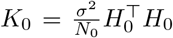, where *N*_0_ is the input dimension. In addition, *V*_0_ and *M*_*l*_ are defined from the eigendecomposition of

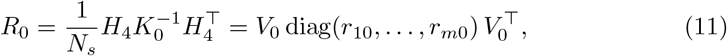

where *N*_*s*_ is the number of training samples (14 stimuli in this paper), *V*_0_ contains the orthonormal eigenvectors of *R*_0_, and {*r*_*k*0_} are the corresponding eigenvalues. The diagonal matrix *M*_*l*_ = diag(*m*_1*l*_, …, *m*_*ml*_) is defined by

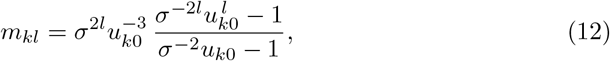

where *u*_*k*0_ is the positive solution of

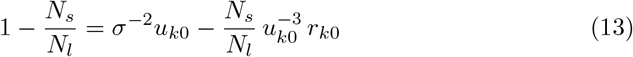

(see [48] for derivation). The correlation matrix for the *l*-th hidden layer is

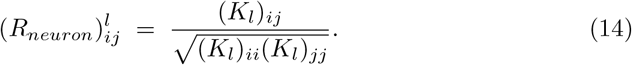

#### 4.4.5 Comparing stimulus similarity between different sample sets

In this paper, stimulus similarity was defined by Pearson correlation coefficient matrices that were computed from either the sensory population responses or the behavior matrix. To assess whether sensory coding aligned with behavioral outputs, we modeled neural activity with a multivariate Gaussian distribution *N*(**0**, *R*_*beh*_), where *R*_*beh*_ was the Pearson correlation coefficient matrix (14 × 14) of the behavior matrix (19 × 14).

If *R*_*beh*_ is full-rank, we can estimate the likelihood of ROI responses through the probability density function of this 14-d Gaussian distribution. Let *C* be the population response matrix (*n*_neuron_ × 14) (Figure 6d, Extended Data Figure 7). We first normalized and centered *C* by computing a z-score for each stimulus:

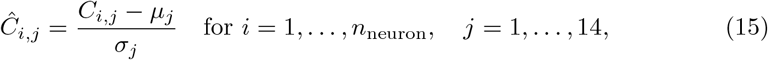

where *µ*_*j*_ and *σ*_*j*_ are the mean and standard deviation of the *j*-th column of *C*. Then the log-likelihood of the 14-d ROI response vector **c**_*i*_ is

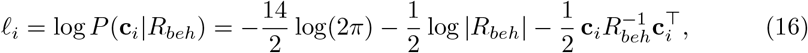

where |*R*_*beh*_| denotes the determinant of *R*_*beh*_, and the mean log-likelihood of the population is

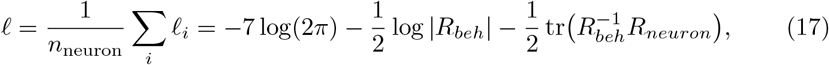

where 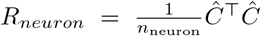 is the Pearson correlation coefficient matrix of the sensory population response.

However, since the dimensionality of behavioral outputs is less than the number of presented stimuli, *R*_*beh*_ is not full-rank. We overcame this singularity by performing likelihood estimation in the subspace spanned by the principal components of *R*_*beh*_,

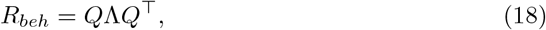

where *Q* is the orthogonal matrix of eigenvectors of *R*_*beh*_ and Λ is the diagonal matrix of eigenvalues of *R*_*beh*_. The eigenvector basis, denoted by *Q*_*i*_ for *i* = 1, · · ·, 14, consists of the columns of the matrix *Q*. We selected the first 7 principal components, denoted by 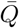, which explained 98.6% of the variance. Then the Gaussian distribution can be defined in this 7-d subspace notated as 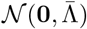, where 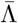 represents the 7 × 7 diagonal matrix of the first 7 eigenvalues of *R*_*beh*_. We then projected *Ĉ* into the 7-d subspace:

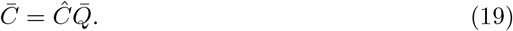

Therefore, the likelihood of each ROI estimated from the Gaussian distribution with behavioral correlation matrix can be computed in this 7-d space. If we notate the *i*-th row in 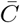 as 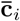, its log-likelihood can be computed by

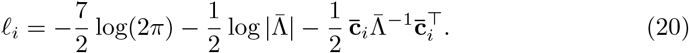

Therefore, the mean log-likelihood over all ROIs in 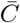 is

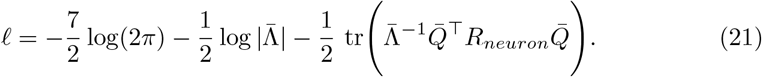

A potential issue is that ROI responses may not lie in the 7-d subspace. As a result, although we normalized each ROI to unit *L*_2_ norm in the original 14-dimensional space, the norm of its projection can vary. In the extreme, 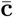 would be zero if the response vector is orthogonal to the subspace. In Eq. 20, this issue can bias its third term, so some ROIs may achieve spuriously high likelihoods because much of their response energy lies outside the 7-d subspace, rather than because they align with the covariance structure of the 7-d Gaussian. On the population level, it is also reflected in Eq. 21, wherein a population exhibit spuriously high likelihood because the entries in 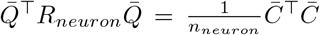 are too small. To enable fair comparisons of stimulus similarity across neural populations, we rescaled the projected responses by a correction factor *γ*:

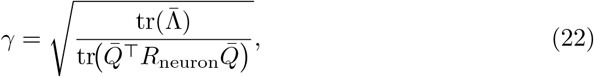

where 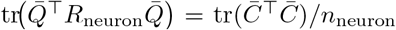 represents the mean squared *L*_2_ norm of neurons in the 7-dimensional subspace. This normalization equalizes the mean *L*_2_ norm in the 7-d space across populations to solve the above projection issue. The corrected log-likelihood of a single ROI is

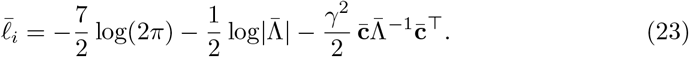

Accordingly, the expected corrected log-likelihood for a population is

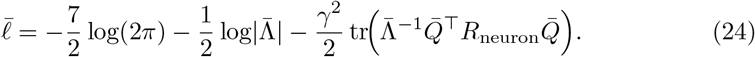

For the simulation in Figure 4, the features of each hidden layer formed a 512 × 14 response matrix. The mean log-likelihood was computed from the individual log-likelihood in Equation 23. For the theoretical prediction in Figure 4, the mean log-likelihood of each layer were computed by Equation 24, with *R*_neuron_ evaluated according to Equation 14.

For likelihood estimation of experimental data in Figure 5, neurons in each brain region across all fish formed the *n*_*neuron*_ × 14 matrix. Then the log-likelihood of each ROI was estimated by Equation 23. We averaged the log-likelihood of neurons from each brain area for each fish; we then averaged these mean log-likelihoods across fish for each brain region. The statistical test across fish in Extended Data Table 1 was based on the region-level mean log-likelihood and used a one-sided Wilcoxon signed-rank test from the SciPy package [84].

#### 4.4.6 Predicting sensory responses by behavioral readout

Given readout weights *U*, sensory responses of 1000 neurons *C*^*M*^ (1000 × 14) were modeled to be linearly transformed into the motor response:

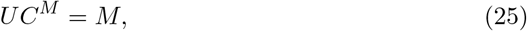

where *M* was the z-scored 19 × 14 behavior matrix for each stimulus (similar to Eq. 15). The readout weights *U* were defined by independent samples from an uncorrelated Gaussian distributed to keep the behavioral alignment [85] (Supplementary Information). Since the dimensionality of *U* is higher than that of *M*, the number of possible solutions for *C*^*M*^ is infinite. We therefore introduced an efficiency constraint to focus on the solution with minimal activity (measured by the *L*_2_ norm). The solution to this linear model of efficient coding of stimuli driving behavior can be found by an optimization problem:

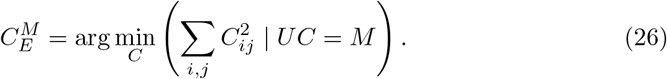

In the Supplementary Information, we analyze this problem and derive a closed-form solution,

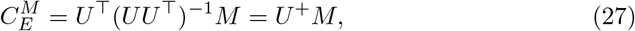

where *U* ^+^ is the pseudo-inverse (or Moore-Penrose inverse) of *U* . When the population size is sufficiently large, the solution can be approximated by

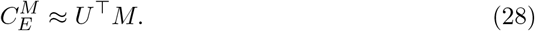

The linear model solution was balanced between negative and positive values, but the recorded sensory responses were mostly positive due to the constraint of non-negative firing rates. Therefore, we added a non-negativity constraint on the solution, which we called the full nonlinear model:

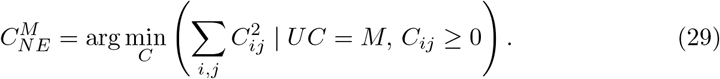

This quadratic programming problem does not have an analytical solution. Instead, in the Supplementary Information we derive the implicit-form solution,

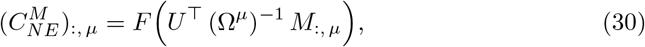

where *F* is the ReLU (rectified linear unit) function, *A*_:,*µ*_ denotes the *µ*-th column of the matrix *A*,

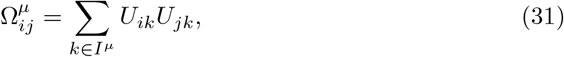

and

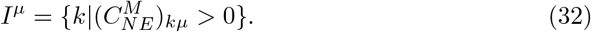

The solution for the full nonlinear model can also be approximated when the population size is sufficiently large,

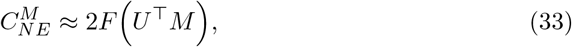

where this approximation aligned very well with the result shown in Fig. 5.

The detailed analytical derivation of those two models can be found in Supplementary Information. For the simulation, we used a Python package, Cooper [86], to solve the constrained optimization problem.

### 4.5 Generative AI

During the preparation of this work the authors used ChatGPT in order to smooth and refine the language. After using this tool/service, the authors reviewed and edited the content as needed and take full responsibility for the content of the published article.

---

**Extended Data Figure 1.**
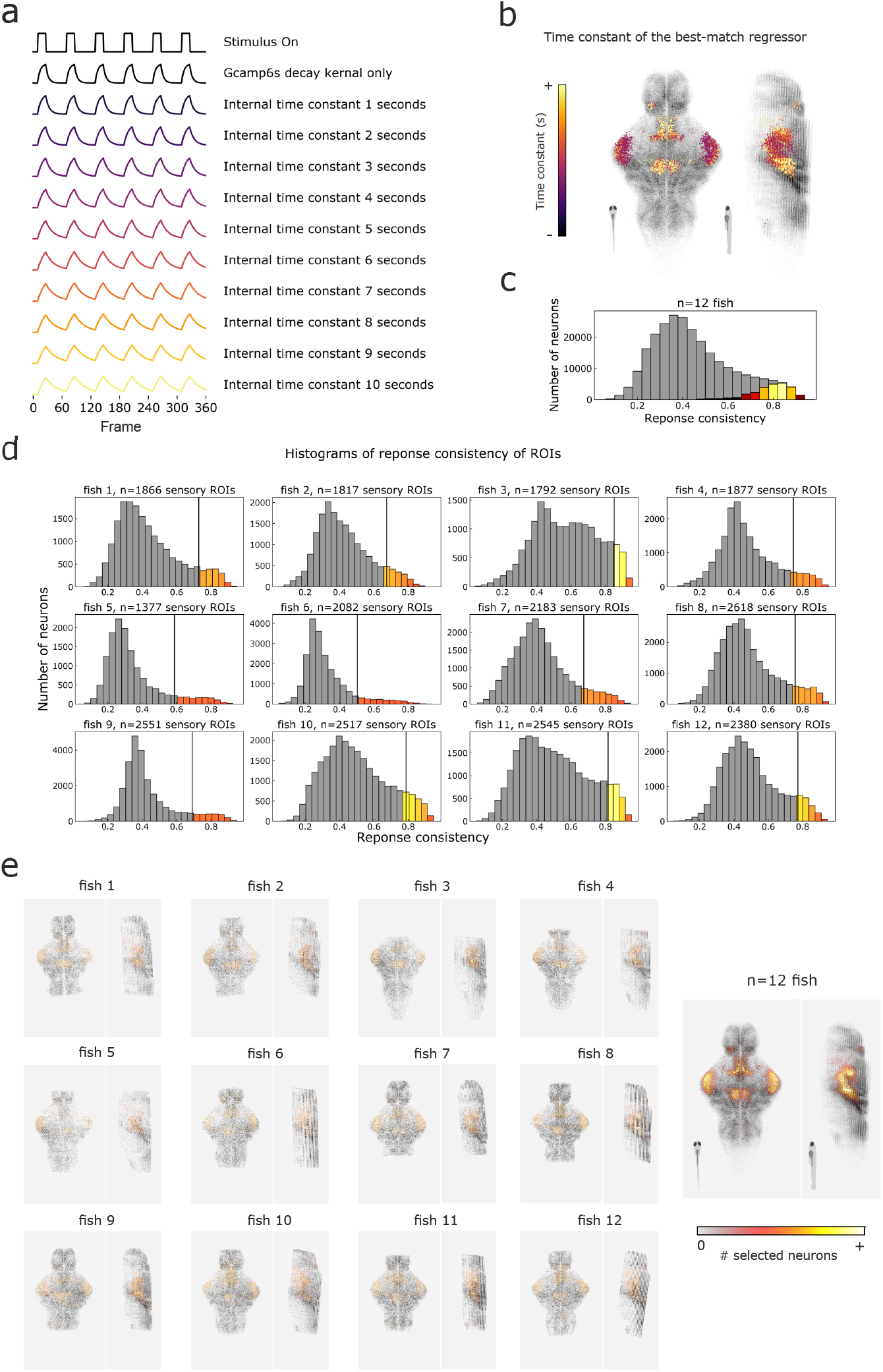
Whole-brain screening of sensory neurons for each fish. **a**, Regressors with different time constants used for computing sensory consistency. **b**, The brain map of time constants of the best-matching regressor for each ROI. **c, d**, Histograms of sensory consistency for all (grey) and sensory (colored) ROIs across 12 fish (**c**) and individual fish (**d**). **e**, Top and side view of the density heatmap of all detected ROIs (grey) and sensory ROIs (colored) across 12 fish and individual fish. Fish are registered onto the reference brain.

**Extended Data Figure 2.**
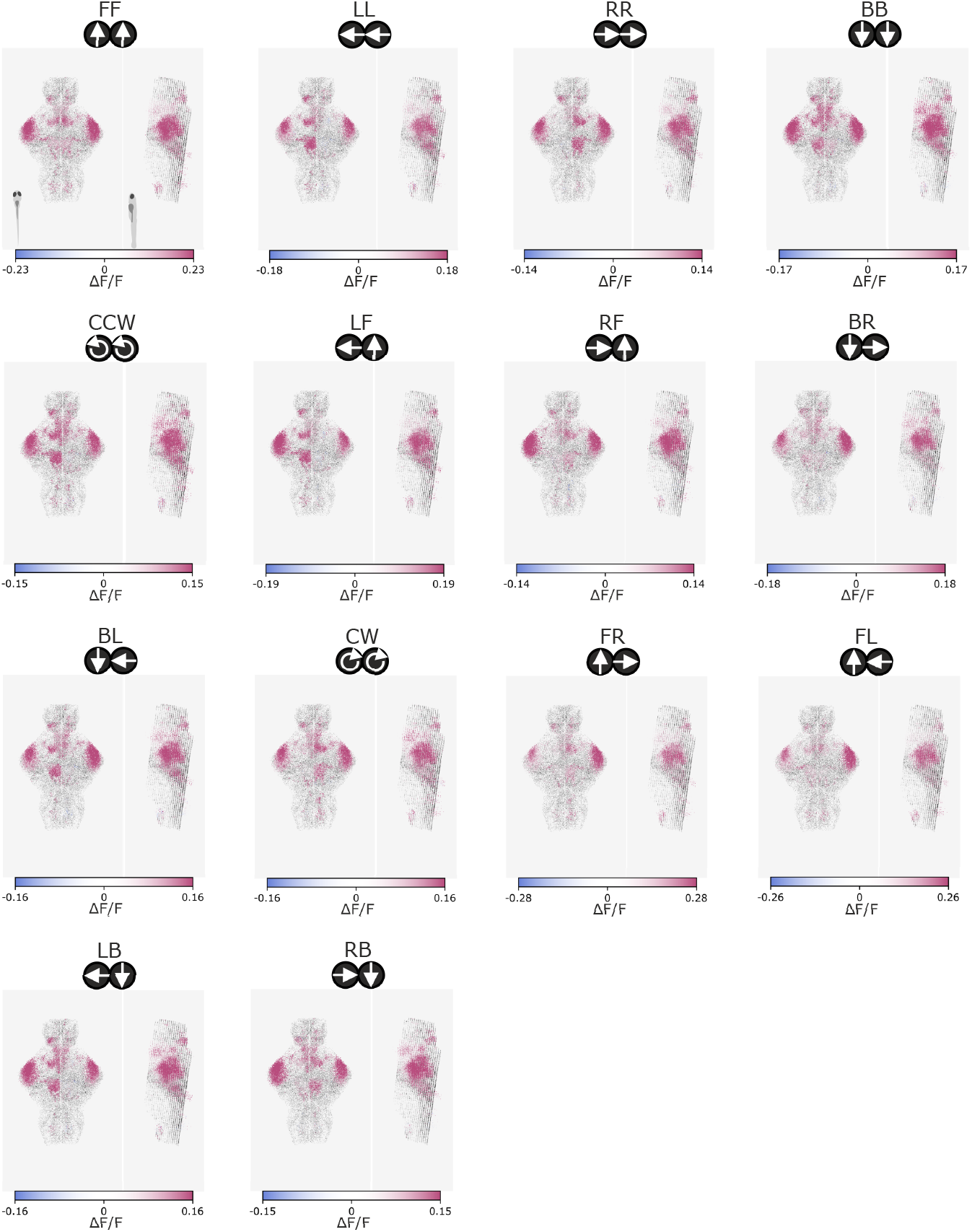
Brain activation map for individual visual stimuli. The heatmap shows the activation of sensory ROIs across 12 fish during motion stimuli. All other detected ROIs are displayed in grey.

**Extended Data Figure 3.**
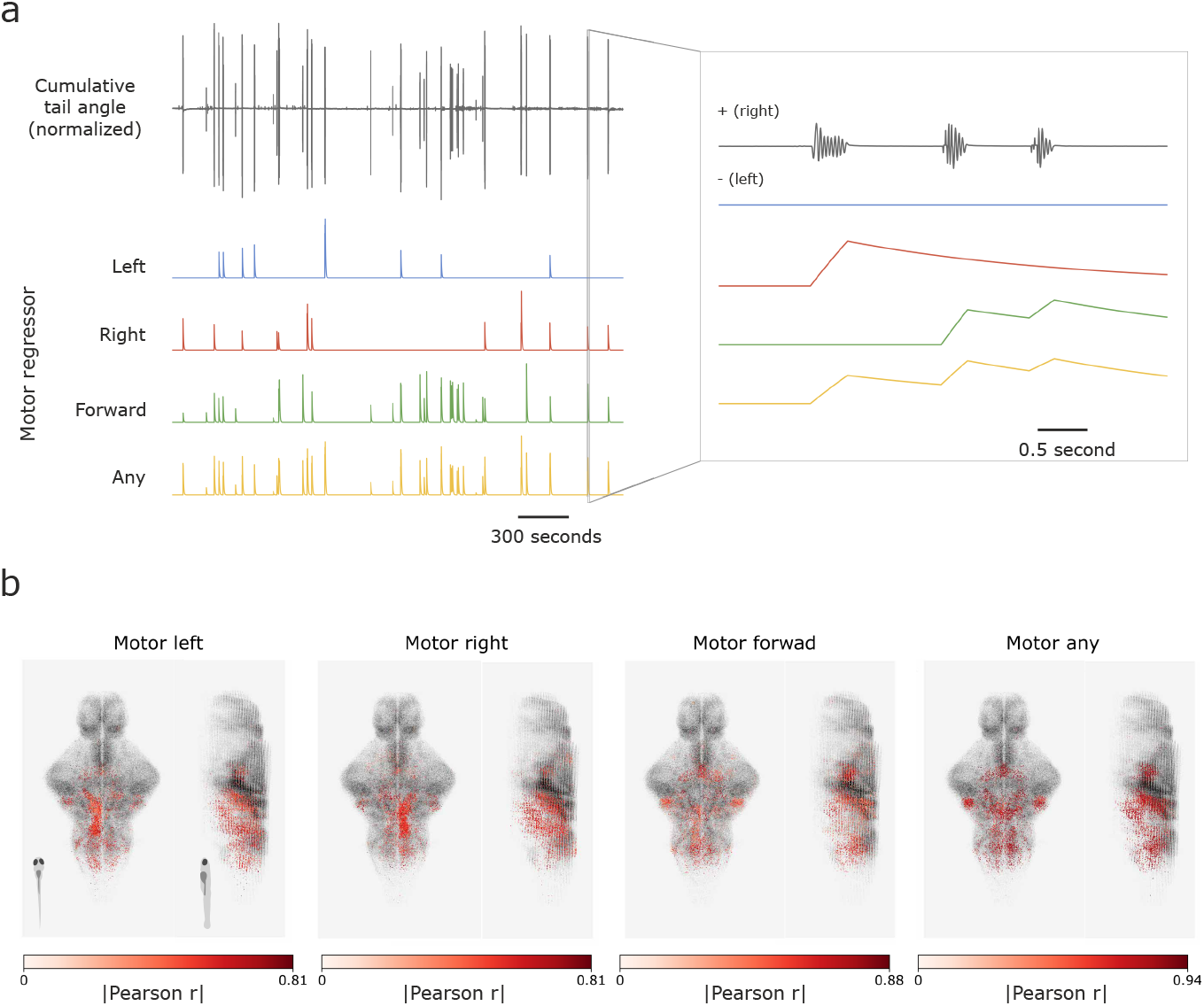
Brain activation map for motor ROIs that correlate with individual bouts. **a**, Left: Example traces of cumulative tail angle (top) and motor regressors (bottom). Right: Zoomed-in traces of tail angle and regressors. **b**, Top view of voxelized heatmap showing the 2% of neurons in each fish with the highest correlations to each motor regressor across 12 fish.

**Extended Data Figure 4.**
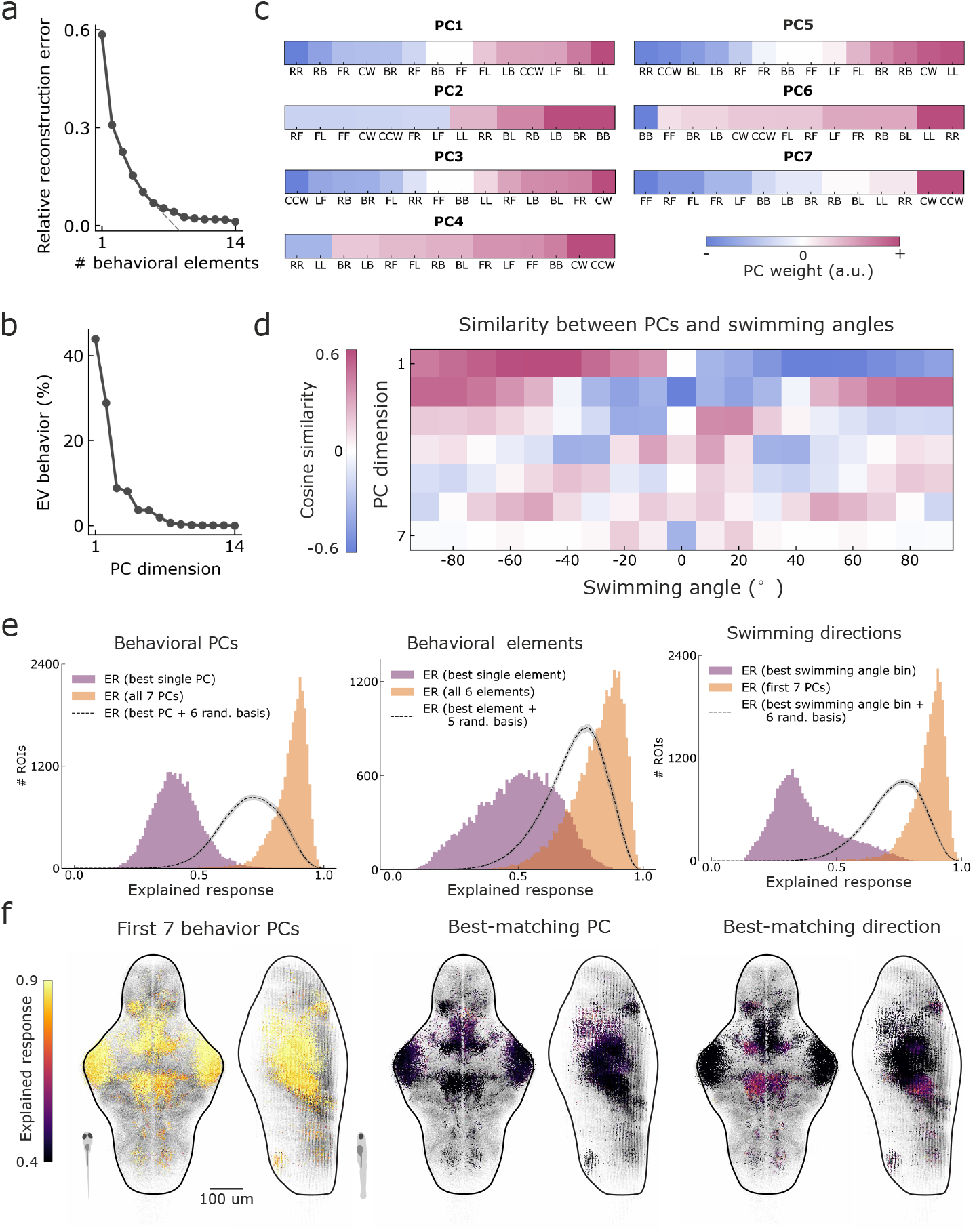
Explaining sensory responses using behavioral principal components. **a**, Relative reconstructive error in sparse dictionary learning for different numbers of behavioral elements. **b**, Explained variance of behaviors by principal component dimensions. **c**, Weights of behavioral principal components for each stimulus. Stimulus order is sorted by its weight in each principal component. **d**, Cosine similarity between each principal component and bout frequencies within each bout angle bin. **e**, Histogram of the explained response of each ROI by principal components, behavioral elements learned by sparse dictionary learning and bout frequencies within each bout angle bin. **f**, Brain maps of explained response for all principal components, best-matching principal component and best-matching angle bin.

**Extended Data Figure 5.**
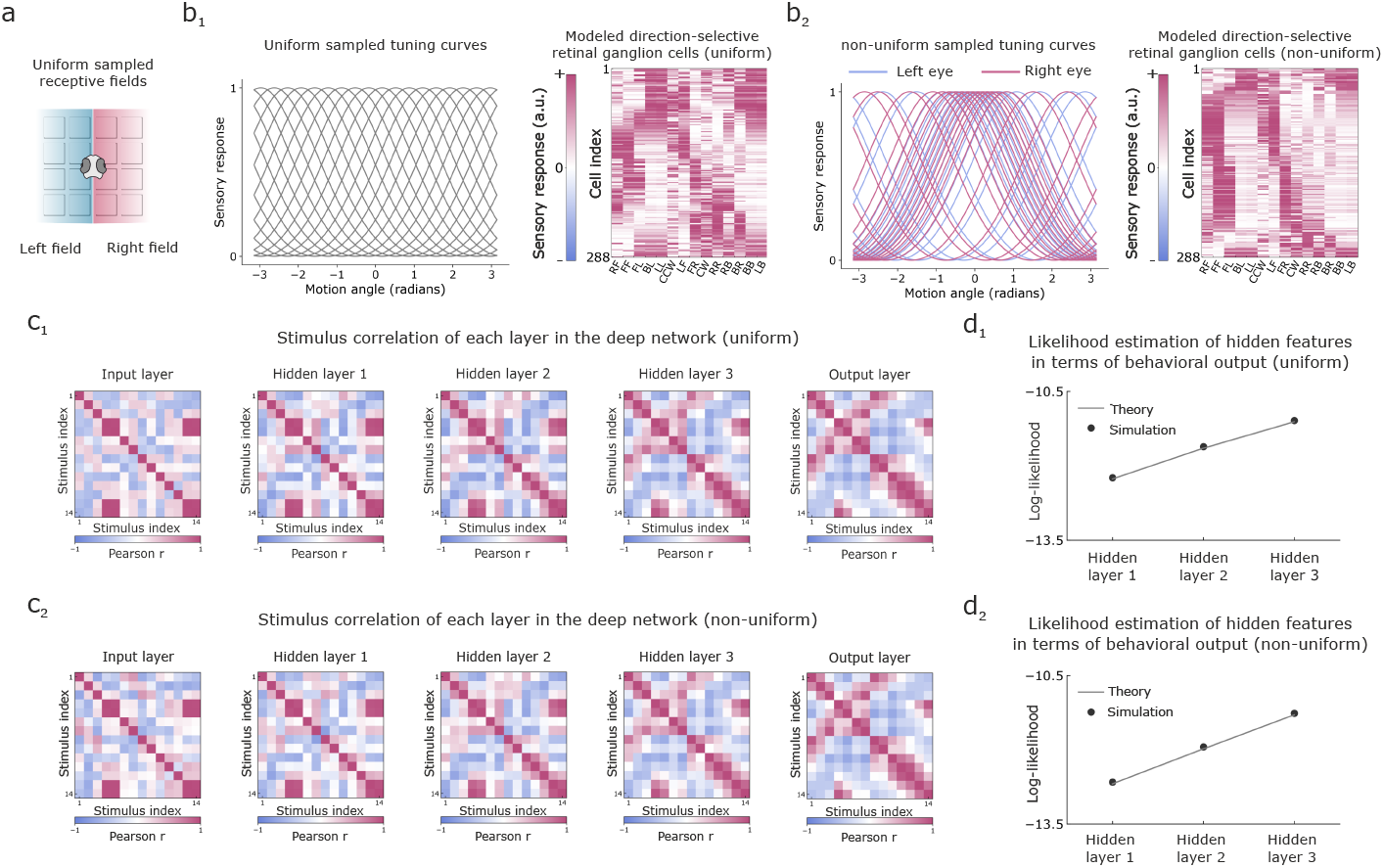
Quantification of sensorimotor transformation in deep linear networks with modeled retinal input. **a**, Receptive fields used in retina modeling. Sixteen receptive fields are uniformly sampled, with eight in each monocular hemifield. **b**, Tuning curves and responses of modeled directional-selective retinal ganglion cells. **b**_**1**_, each receptive field contains 18 neurons uniformly tuned from -pi to pi. **b**_**2**_, each receptive field contains 18 neurons tuned to the directions based on Kramer et. al., 2019. **c**, Stimulus similarity matrix of each layer in the deep linear network trained by uniformly tuned retinal input (**c**_**1**_) and non-uniformly tuned retinal input (**c**_**2**_). **d**, Likelihood estimation of features in hidden layers in terms of stimulus similarity matrix of behavioral output for uniformly tuned retinal input (**d**_**1**_) and non-uniformly tuned retinal input (**d**_**2**_).

**Extended Data Figure 6.**
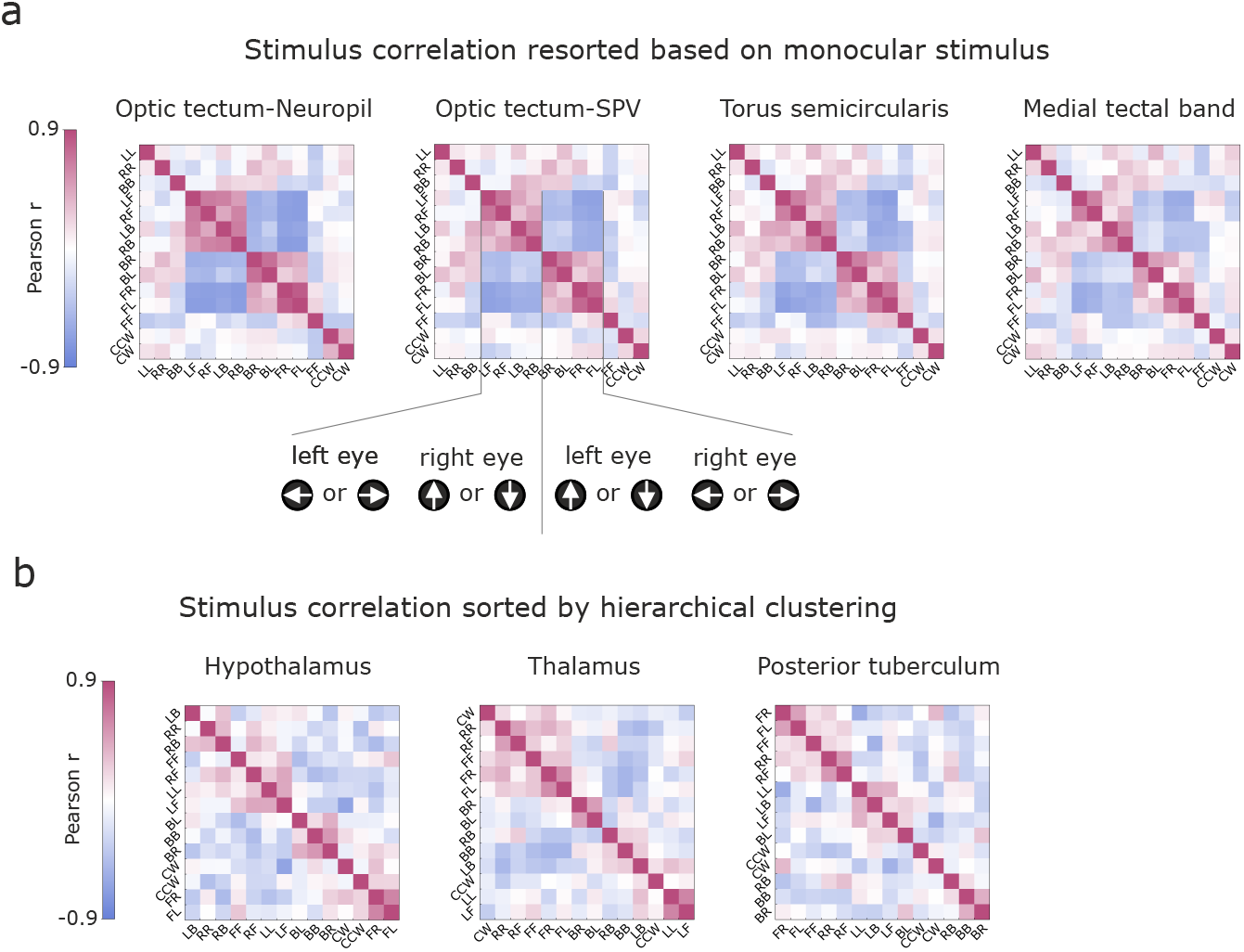
Stimulus similarity in other brain regions. **a**, Stimulus similarity of four brain regions that are strongly shaped by monocular stimulus. **b**, Stimulus similarity of three brain regions that do not exhibit obvious interpretable patterns.

**Extended Data Figure 7.**
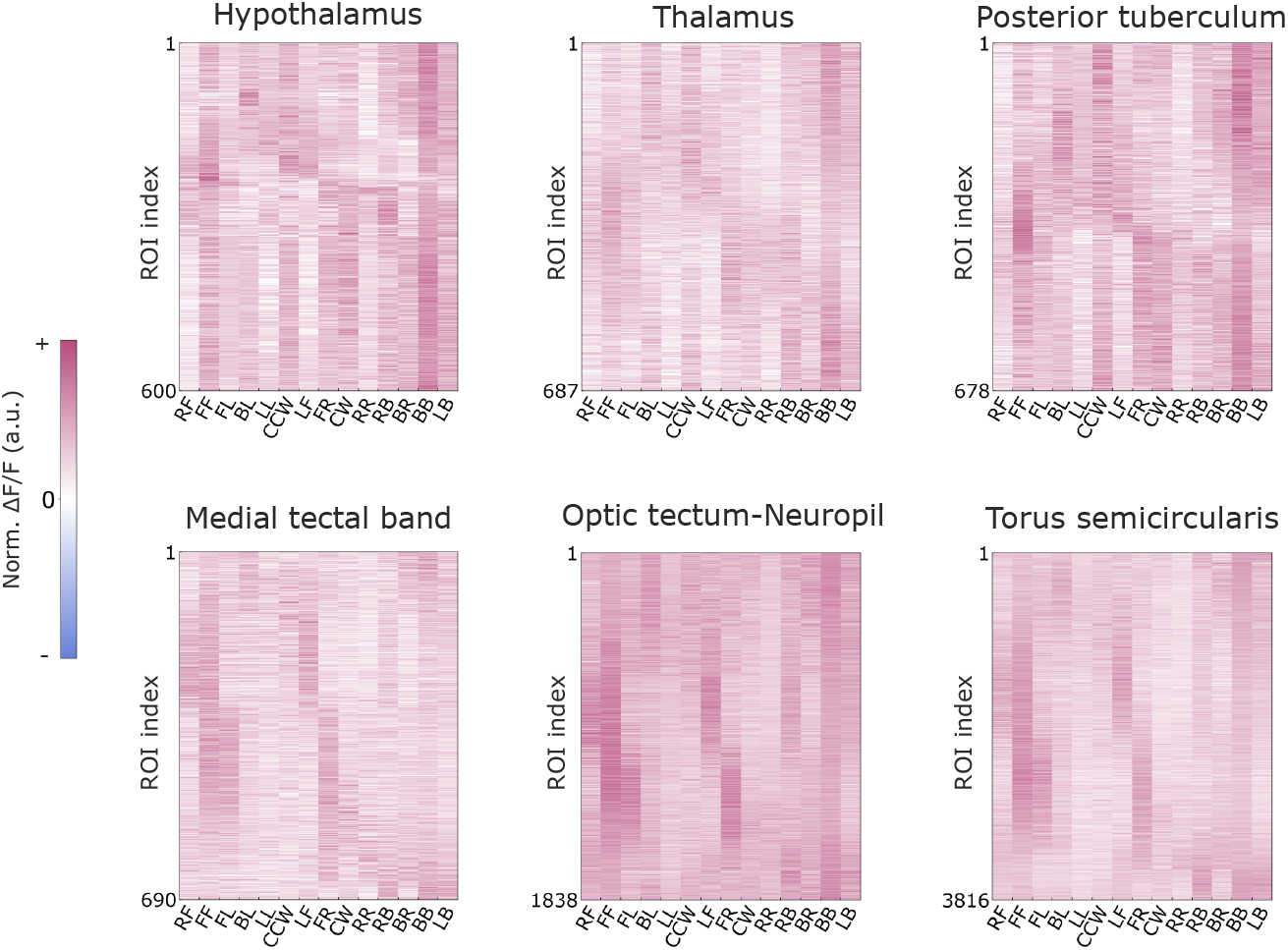
Neural activities of other brain regions. Δ*F/F* of selected sensory ROIs under 14 stimuli in six brain regions. Neural activities are normalized for each ROI.

**Extended Data Figure 8.**
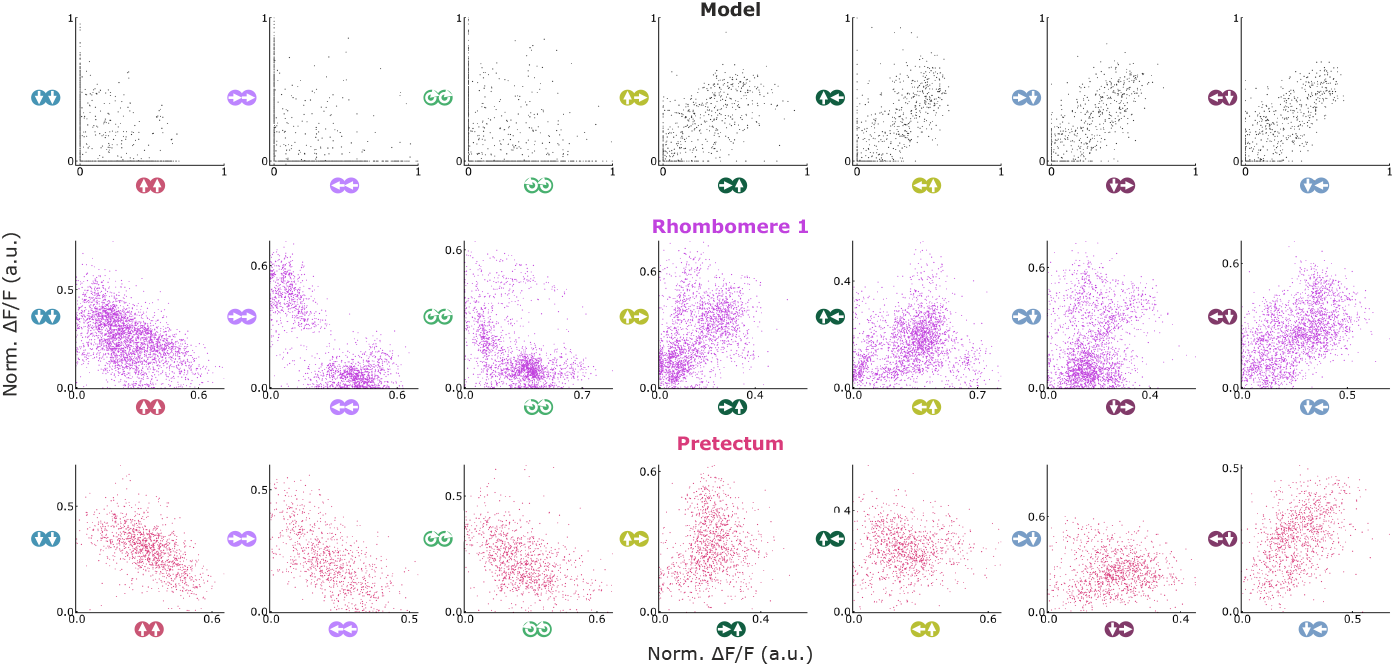
Comparison between the modeled sensory activity and experimental data. The scatter plot shows individual neurons responding to pairs of stimuli. The full nonlinear model captures the correlation between pairs of stimuli, and the neuronal activity in Rhombomere 1 is more aligned with the model compared to that in pretectum.

**Extended Data Figure 9.**
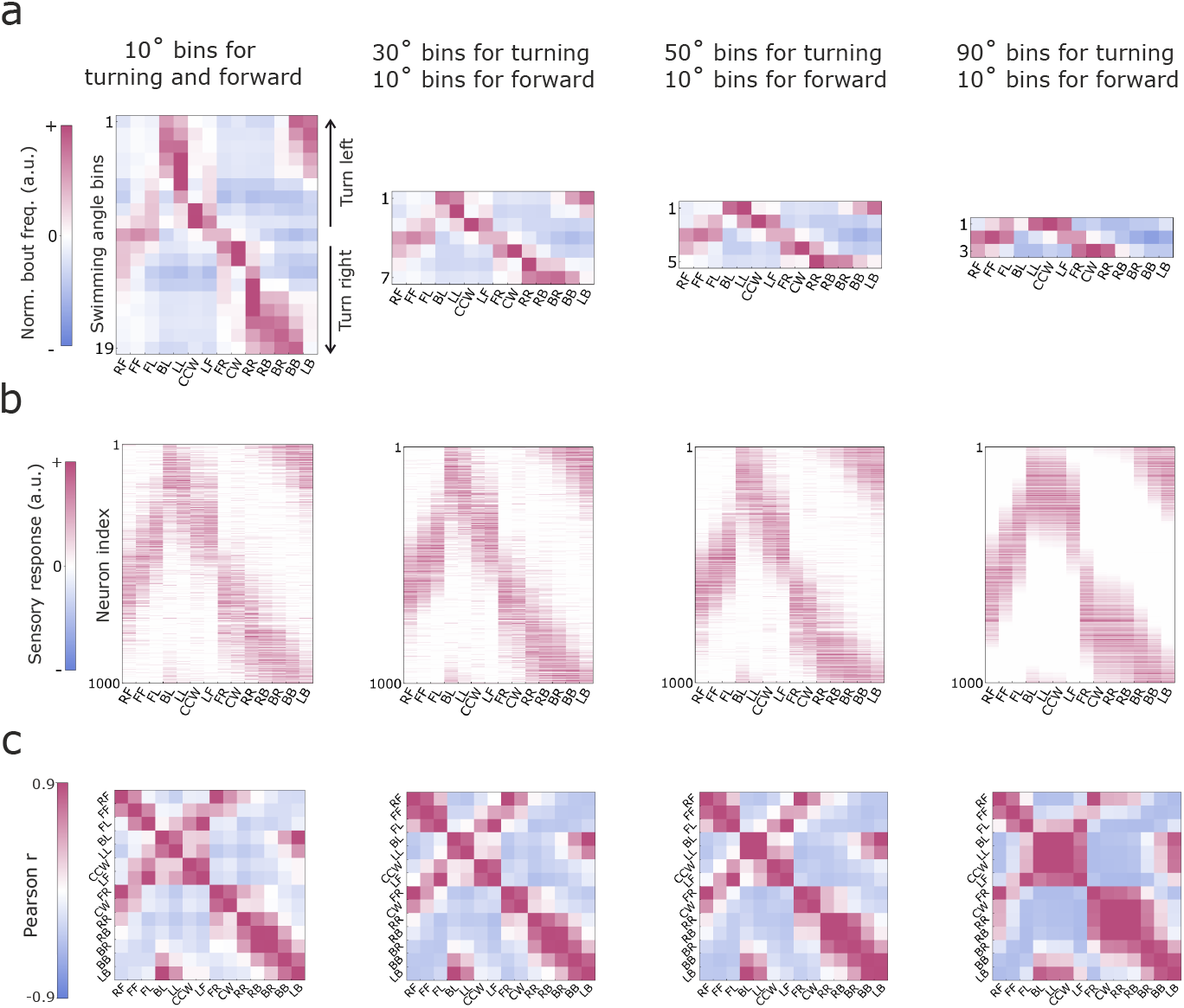
modeled sensory activity at different levels of behavioral granularity. **a**, Behavioral quantification at different granularity. **b**, Sensory activity and **c**, Stimulus similarity matrix from the full nonlinear model under different behavioral quantification. When the behavioral quantification is too coarse-grained, stimulus similarity is distorted so that the sensory activity cannot be correctly modeled from behavior.

**Extended Data Figure 10.**
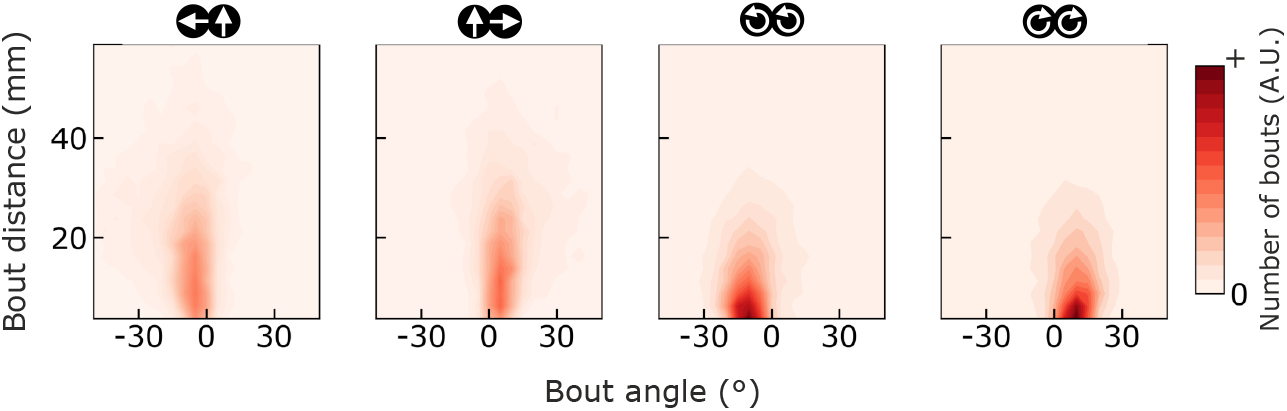
Fine behavioral quantification increases stimulus discriminability. The heatmap shows bout distance and bout direction for observed bouts under four different stimuli. Although the direction of bouts are similar, the difference in bout distance may further distinguish the behavioral responses to different stimuli.

**Extended Data Table 1:** Comparison of behavioral alignment across different brain regions. p-values from one-sided Wilcoxon signed-rank tests comparing the mean estimated log-likelihood values for each pair of brain regions (n = 10-12 fish).

|  | Rh2 | Tg | Pr | Hb | pTu | Th | TS | OT-SPV | hTh | OT-MB | OT-Np |
| --- | --- | --- | --- | --- | --- | --- | --- | --- | --- | --- | --- |
| Rh1 | **0.0046** | **0.0061** | **0.0461** | **0.0093** | **0.0005** | **0.0005** | **0.0002** | **0.0002** | **0.0002** | **0.0002** | **0.0002** |
| Rh2 |  | 0.1697 | 0.311 | 0.1602 | **0.0081** | **0.0024** | **0.0002** | **0.0002** | **0.0005** | **0.0002** | **0.0002** |
| Tg |  |  | 0.4548 | 0.4155 | **0.0017** | **0.0012** | **0.0002** | **0.0002** | **0.0005** | **0.0002** | **0.0002** |
| Pr |  |  |  | 0.4155 | **0.0012** | **0.0012** | **0.0005** | **0.0005** | **0.0017** | **0.0002** | **0.0005** |
| Hb |  |  |  |  | 0.0737 | **0.0161** | **0.001** | **0.0005** | **0.0034** | **0.0005** | **0.0005** |
| pTu |  |  |  |  |  | 0.3667 | 0.1506 | 0.0757 | **0.0134** | **0.0081** | **0.0261** |
| Th |  |  |  |  |  |  | 0.2349 | 0.0757 | **0.0386** | **0.032** | **0.0171** |
| TS |  |  |  |  |  |  |  | 0.1167 | 0.2349 | 0.1506 | **0.0386** |
| OT-SPV |  |  |  |  |  |  |  |  | 0.4849 | 0.2119 | 0.1167 |
| hTh |  |  |  |  |  |  |  |  |  | 0.5151 | 0.5452 |
| OT-MB |  |  |  |  |  |  |  |  |  |  | 0.6333 |

## Supplementary information

## 1 Mathematical models of efficient coding of stimuli driving behavior

### 1.1 Closed-form solution of the linear model

The linear model is defined by

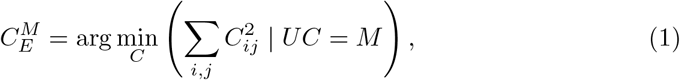

where 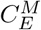 is the optimal solution, *U* are the readout weights, and *M* represents the matrix of motor outputs. The dimensionalities of these matrices are denoted as 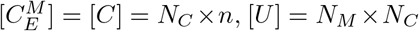, and [*M*] = *N*_*M*_ ×*n*, where *N*_*C*_ is the number of neurons in the population, *N*_*M*_ is the number of motor elements, and *n* is the number of stimulus conditions. For simplicity, we assume that *U* and *M* are both full rank. Since the number of neurons in a brain area typically exceeds the number of motor elements, we also assume that *N*_*M*_ ≤ *N*_*C*_.

We introduce a Lagrange multiplier matrix Λ for the constraint. Define the Lagrangian:

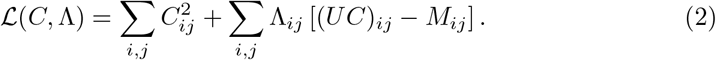

In a matrix form, it can be written as

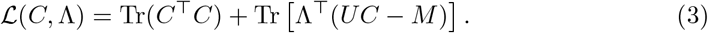

Taking the derivative of *L*(*C*, Λ) with respect to *C* gives

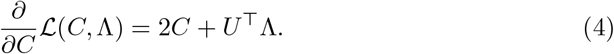

Setting the derivative equal to zero, we find that the optimal solution satisfies

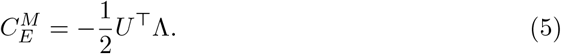

Plugging this expression back into *UC* = *M*, we get

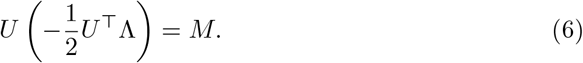

Therefore,

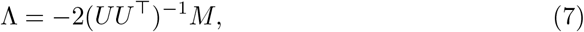

where the matrix inverse is well-defined because the rank of *U* is *N*_*M*_ by assumption. Substituting back, we find the solution:

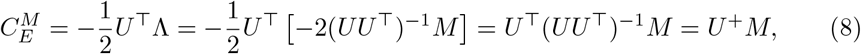

where *U* ^+^ is the pseudo-inverse (or Moore-Penrose inverse) of *U* . The modeled neural response *C*^*M*^ are effectively “reverse projected” from motor outputs to the space of neural responses.

### 1.2 Implicit solution of the full nonlinear model

The full nonlinear model is defined by the following quadratic programming problem:

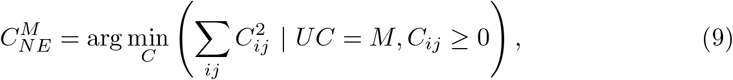

where 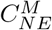 is the optimal solution of the model, notated as *C*^∗^ for simplicity, and all other variables are defined as above.

This is a minimization problem subject to both equality and inequality constraints. The optimization function to be minimized is

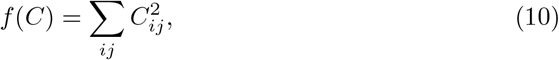

the inequality constraints are

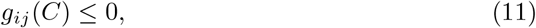

where

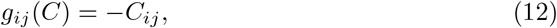

and the equality constraints are

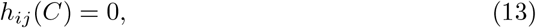

where

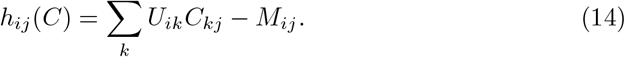

For *C*^∗^ to be a local minimum, the stationarity component of the Karush-Kuhn-Tucker (KKT) conditions [1, 2] implies that there exist coefficients *µ*_*ij*_ and *λ*_*ij*_ such that

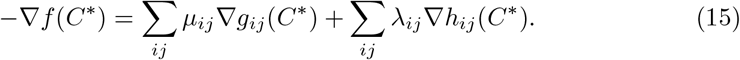

The derivatives for our problem are defined component-wise by

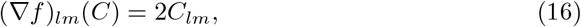

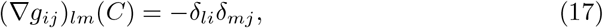

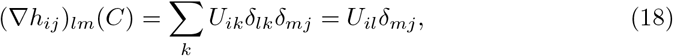

where *δ*_*ij*_ denotes the Kronecker delta function, which is defined to be 1 if its indices are equal and 0 otherwise. Therefore, the stationarity condition says

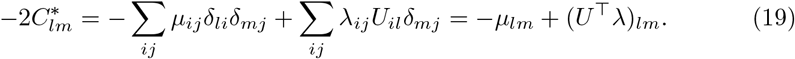

In matrix notation, this is

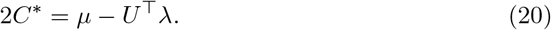

The complementary slackness component of the KKT conditions says

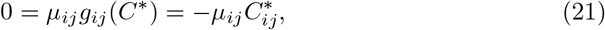

which implies that either 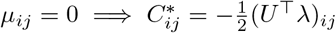 or 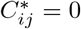. Primal feasibility guarantees that 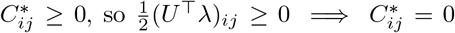. On the other hand, dual feasibility guarantees that 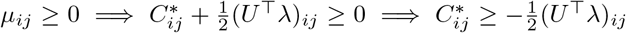. Therefore 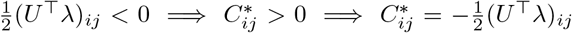. Putting these pieces together, we have shown that

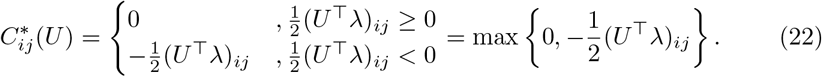

Put differently, we see that

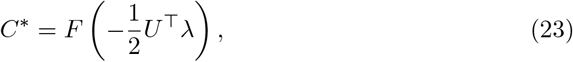

where *F* is the ReLU (rectified linear unit) function. Plugging this into the stationarity condition, we’ve also shown that

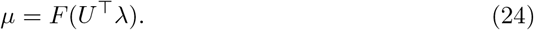

These expressions for *C*^∗^ and *µ* are enough to guarantee stationarity, primal feasibility of the inequality conditions, dual feasibility, and complementary slackness.

The only remaining component of the KKT conditions is the primal feasibility of the equality constraints:

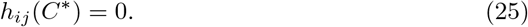

Consequently, the matrix equation

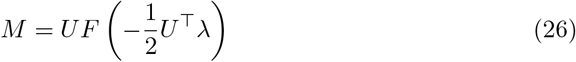

implicitly defines the values of Lagrange multipliers as a function of *U* and *M* . If not for the nonlinearity (i.e., if *F* (*x*) = *x*), the solution would be the simple expression we derived in the previous section (Eq. 7). A similar form implicitly specifies the Lagrange multipliers in the presence of the thresholding nonlinearity. In particular, note that we can account for the nonlinearity by writing

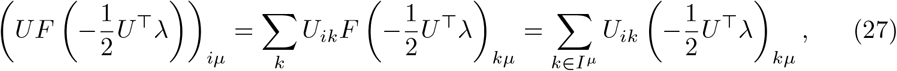

where

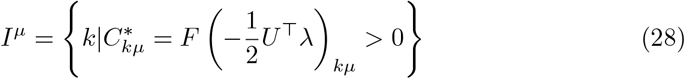

is the set of neurons activated in the *µ*-th stimulus condition. If we then define

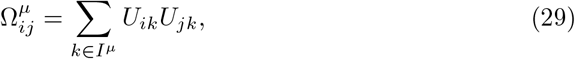

then the above equation simplifies to

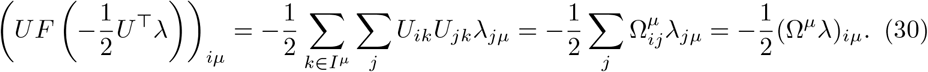

We therefore find that the Lagrange multipliers are

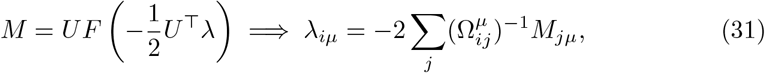

where we’ve assumed that the *N*_*M*_ × *N*_*M*_ matrix Ω^*µ*^ is full rank. This is reasonable to expect if *N*_*C*_ ≫ *N*_*M*_, and the sparsity of *C*^∗^ is not too high. We’ve therefore found that

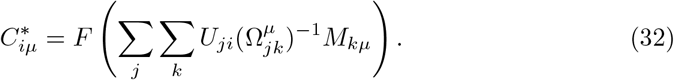

Although this is an exact expression for *C*^∗^, the circular dependence of *I*^*µ*^ on *M* and *U* limits its practical utility to special circumstances. For instance, we’ll see below how we can understand its form when *U* is a random matrix. In a matrix notation, the solution can be written as

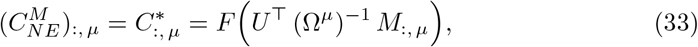

where *A*_:,*µ*_ denotes the *µ*-th column of the matrix *A*.

## 2 Statistical arguments

We have defined the connection weights *U* as samples independently drawn from a Gaussian distribution *N*(0, 1*/N*_*C*_). Here we derive several important consequences of this assumption.

### 2.1 Preserved similarity structure

For a constant z-scored matrix 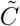 with same shape of *C*, the expectation of the Pearson correlation between the *i*-th and *j*-th columns in 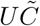 is the same as the Pearson correlation between the *i*-th and *j*-th columns in 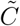. To see this, first note that

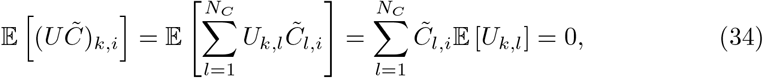

where E denotes the expectation of elements of *U* over the *N*(0, 1*/N*_*C*_). The covariance between the *i*-th and *j*-th columns in 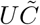 is thus

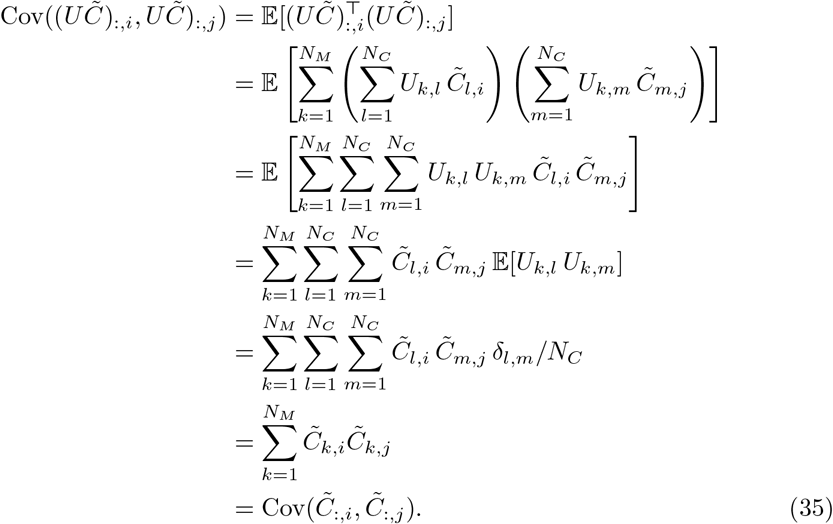

Note that since the matrix 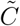 is z-scored by assumption, this implies that the variance of each column of 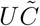 is one. Therefore, the columns of 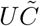 and 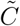 are zero mean and unit variance, and the Pearson correlation matrices, *e*.*g*.

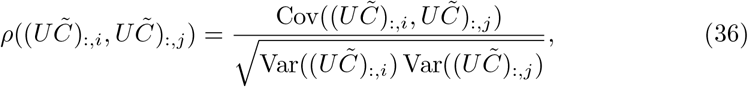

are the same as the covariance matrices. This implies that

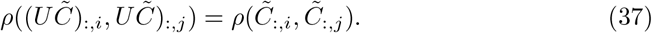

In other words, this Gaussian projection preserves the pair-wise stimulus similarity of 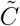.

### 2.2 Statistical properties of the two solutions

We next use the randomness of *U* to derive simpler expressions for the solutions of the linear and full models when *N*_*C*_ is large.

For the linear model, recall that

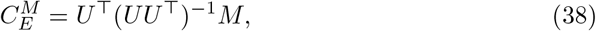

and consider the matrix Ω = *UU* ^⊤^ that enters this expression. Its expected value is straightforward to evaluate:

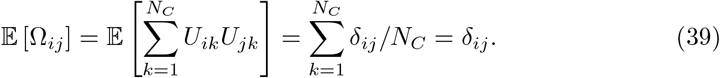

Therefore, E[Ω] is the identity matrix, 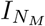. For large values of *N*_*C*_, Ω does not vary much from its expected value. To see this, we first apply Wick’s theorem to evaluate the second moments of Ω:

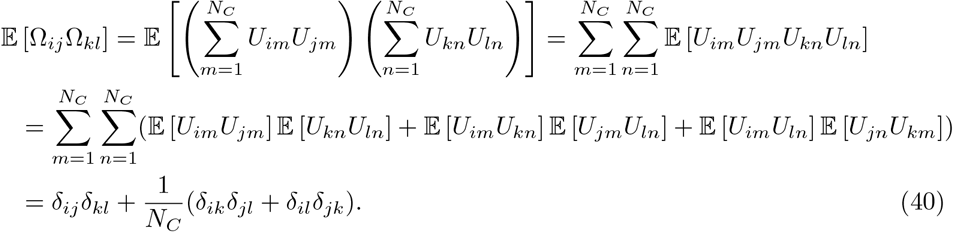

The covariance is accordingly,

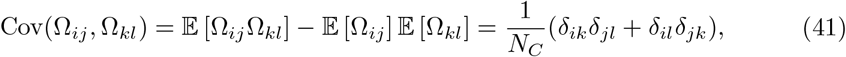

which goes to zero as *N*_*C*_ gets large. The matrix Ω thus approaches the identity matrix as *N*_*C*_ increases, with convergence characterized by a rate proportional to 1*/N*_*C*_. As a result,

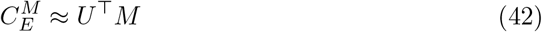

when *N*_*C*_ is sufficiently large. Conceptually, the pseudo-inverse of *U* approaches *U* ^⊤^ in this limit because rows of *U* are statistically uncorrelated and unit norm.

For the full nonlinear model, recall that the solution is

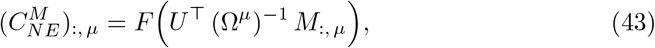

where

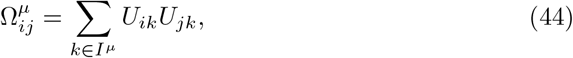

and

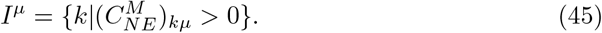

The matrix Ω^*µ*^ can be analyzed similarly to Ω, except now the sums are limited to *I*^*µ*^. Therefore, for large values of *N*_*C*_

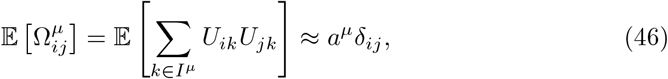

where *a*^*µ*^ = 0.5 is the expected fraction of neurons active in pattern *µ*, and

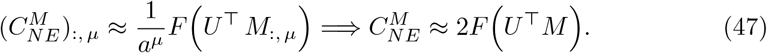

In practice, this approximation agrees extremely well with the exact solutions plotted in Fig. 5. The model essentially reverse projects *M* with *U*^⊤^, rectifies the result to preserve non-negativity, and rescales the final result to compensate for the zeros. In this way, the Pearson correlation coefficients between pair-wise stimuli in 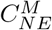 are distorted by the nonlinearity *F*, but they still display patterns similar to that of *M*.

